# Cryoaerosolization Enables Scalable Vitrification-Based Cell Cryopreservation

**DOI:** 10.64898/2026.07.24.740597

**Authors:** Joseph R. Kangas, Abhilash Ojha, Minhan Jiang, Mohammad Shameem, Bhairab N. Singh, John C. Bischof, Christopher J. Hogan

## Abstract

Cell therapies hold transformative potential for treating cancer, neurologic disorders, organ failure, diabetes, and other conditions, but their widespread clinical deployment is constrained by the lack of scalable cryopreservation methods that maintain high post-thaw viability. Current standard practice using slow freezing can lead to cell death and impaired cell function. Vitrification offers an alternative by cooling samples rapidly enough to bypass ice formation entirely, better preserving cell structure and function. The high cooling and warming rates required for vitrification have previously been achieved by quenching microliter-scale samples directly into convective cooling and warming baths. Here, we present a cryopreservation platform that overcomes the throughput limitations of existing systems by combining a vibrating orifice aerosol generator with an impinging conical nozzle to generate and confine micrometer-scale droplets mid-flight in liquid nitrogen. This approach mitigates cooling losses due to the inverse Leidenfrost effect, increasing cooling and warming rates by nearly an order of magnitude compared to conventional droplet vitrification, while improving throughput by two orders of magnitude. To test the efficacy of this system, we cryoaerosolized and rewarmed human induced pluripotent stem cells, porcine red blood cells, and human dermal fibroblasts using only 190–25 wt% (2.5–3.7 M) permeating cryoprotective agent, achieving >90% post-thaw viability for HDFs and hiPSCs and 94% recovery for RBCs, with retained colony-forming capacity additionally demonstrated in hiPSCs. This work demonstrates the first scalable vitrification-based cryopreservation method capable of achieving both high cooling (≈ 200, 000 K min^−1^) and warming rates (≈ 1, 000, 000 K min^−1^) while maintaining the high-throughput processing required (≥100 mL h^−1^) for next-generation cell therapies.

**Significance Statement:** Cell therapies require robust long-term storage methods to enable widespread clinical deployment. Current approaches utilizing refrigeration or small-scale vitrification cannot meet the scalability and viability requirements for next-generation therapeutics. In this work we demonstrate a cryoaerosolization process that achieves both ultra-rapid cooling rates (>200,000 °C min^−1^) and high throughput (>100 mL h^−1^) by generating micrometer-scale droplets and spraying in a liquid nitrogen impingement stream. Using only as little as 19 wt% cryoprotectant, we achieved >90% cell viability and maintained function, comparable to low-throughput methods, but at two orders of magnitude higher processing rates. We also introduce a just-in-time CPA loading approach that reduces toxicity exposure. This technique enables scalable vitrification-based cryopreservation of large-volume cell products.

---

**A**dvances in cell therapies and regenerative medicine offer an unprecedented opportunity to transform treatments for cancer, neurologic disorders, organ failure, diabetes, and many additional diseases and disorders.(1, 2) These therapies utilize a diverse range of cells, including engineered immune cells (CAR-T, TCR-T, NK cells), stem cells and stem cell–derived populations (HSCs, MSCs, iPSCs), and primary cell therapies such as hepatocytes and islet cells. Most cell-based therapeutics are not manufactured at the point-of-care, necessitating reliable long-term storage methods such as cryopreservation to enable efficient transport and off-the-shelf availability. (3, 4) Recent clinical studies report immune-cell infusion volumes ranging from 20 to 200 mL, while administered doses can range from approximately 10^8^ to 5 × 10^9^ cells, depending on the cell product and indication.(5– 7) Achieving timely clinical availability requires manufacturing platforms capable of generating and preparing therapeutic-scale cell numbers for storage; a recent GMP-grade CAR-T manufacturing study achieved 150–200-fold expansion over 7 days, producing up to 2.6 × 10^9^ cells with > 93% viability before automated formulation for cryopreservation.(8) High-fidelity (>90%) recovery after cryopreservation is key, as dead or damaged cells not only dilute the effective therapeutic dose but may also compromise product safety by provoking unwanted inflammatory responses.(3, 9) Additionally, the need for robust cryopreservation is essential for therapies delivered in multiple doses over weeks to months, not only to preserve product viability, but also to reduce the costs and logistical strain of repeated manufacturing and distribution.(3, 4) Cryopreservation involves cooling biological material to cryogenic temperatures below − 150 ◦C, where the processes normally associated with life (active and passive transport, metabolic activity) effectively cease, stopping the so-called “biological time”(10). The alternatives to cryopreservation are far from ideal; storage under refrigeration leads to gradual loss of viability, while extended in vitro culture requires continuous proliferation that drives unwanted differentiation and genetic drift.(11–14) By maintaining cells in a cryogenic state, both outcomes are avoided, preserving therapeutic potential for decades.(15) The main risk associated with cryogenic storage is that once biological material is cooled below its melting point, it becomes susceptible to ice formation, which can trigger a cascade of damaging effects across multiple size domains. Cell-scale effects of ice formation include changes in pH, osmotic pressure, water availability, cell shape and volume, and membrane integrity; while tissue-scale effects include the destruction of tissue-scale structure such as vasculature and extracellular matrices, either directly from ice formation or from cracking induced by thermal expansion of ice(16–19).

To mitigate ice-induced damage, specific ice-suppressing chemicals called cryoprotective agents (CPAs) are introduced prior to cooling. CPAs suppress ice formation through a variety of mechanisms, modulating the system viscosity, melting and glass transition temperatures, crystal growth rate, and the free energy of nucleation.(20–23) Both the effectiveness (ice-suppressing quality) and chemical toxicity of a CPA are highly dependent on concentration, with higher concentration CPAs more toxic while also showing stronger potential for reducing ice formation. CPAs can be separated into two categories, either penetrating or nonpenetrating, based on whether or not they permeate the cell membrane.(24) Penetrating cryoprotectants readily migrate across cell membranes, directly reducing damaging intracellular ice formation during cooling and warming. Common penetrating CPAs include DMSO, propylene glycol (PG), ethylene glycol (EG), glycerol, ethanol, and methanol. Nonpenetrating cryoprotectants (which are often larger sugars) are typically added in conjunction with penetrating CPAs. These increase the solute concentration in the extracellular space, causing water to leave the cell via osmosis, and hence decrease the freezable fraction of the interior of the cell.

Slow freezing is a cryopreservation method that avoids the particularly damaging intracellular ice formation that can form within cells. It involves placing cells equilibrated in a 10% (w/v) DMSO solution into a cryovial, then cooling the cryovial 1 °C min^−1^ to at least -80 °C.(4) During this process, heterogeneous nucleation sites on the inner surface of the cryovial cause water to crystallize at higher temperatures than within the cells. The reduction in free liquid water then increases the concentration of DMSO in the supercooled fraction, which then drives more DMSO into the cells. If cooling rates are sufficiently slow, ice formation can be entirely relegated to the extracellular space, thus avoiding the most damaging intracellular ice formation. Once the cryovials reach -80 °C intracellular DMSO concentrations may exceed 55% (w/v), which is sufficient to place directly into liquid nitrogen for long-term storage without further crystallization. Rewarming and recovery are carried out via convective warming in a warm water bath, where the high intracellular CPA concentrations again prevent further crystallization. Slow freezing is the most widely adopted cryopreservation method due to its simplicity and repeatability, but it has several key drawbacks. Sample sizes are limited to cryovials approximately 1-5 mL in volume, as it becomes difficult to control extracellular ice formation outside this range. For larger doses of cell immunotherapy this can mean having to rewarm and recover multiple samples for a single dose.(30, 31) Ice formation in the extracellular space can lead to mechanical damage to cells as they are confined between growing ice crystals, leading to cell damage and death, particularly problematic in cell aggregate systems. Long-term exposures to the high concentrations of CPA that occur during slow freezing also can have a variety of toxic effects on the cell, ranging from decreased or altered cell function to cell death.

Conversely, vitrification involves avoiding ice formation entirely, both intracellularly and extracellularly, during cooling and rewarming. This approach involves achieving and storing an amorphous glass state. This glassy state is ideal for long-term storage of biological specimens as metabolic processes cease to function, with the only storage-related damage to cells from electromagnetic radiation, particularly high-energy cosmic rays(32). Vitrification can be achieved by cooling a sample loaded with CPA to its glass transition temperature at a cooling rate faster than its critical cooling rate (CCR). Similarly, on warming, to avoid ice formation the sample must be warmed faster than its critical warming rate (CWR). Both the glass forming tendency (CCR and CWR) and the toxicity of CPAs depend exponentially on CPA concentration, thus low concentration CPAs exhibit low toxicity but require rapid cooling and warming rates to avoid ice formation(29, 33–35).

Vitrification-based approaches to cell cryopreservation using low CPA concentrations have been shown to be superior to slow freezing approaches in many sensitive cell types; however, they rely on rapid quenching of very small volumes (*µL*-scale droplets and straws), and hence lack the scalability required for clinical application.(28, 36–38) Larger-droplet vitrification and mL-scale container-based vitrification methods increase the volume processed per operation, but generally do so at the cost of lower cooling and warming rates and therefore require higher CPA concentrations. In comparison, clinically established slow-freezing methods routinely process cell suspensions in cryobags with capacities ranging from 10 to more than 100 mL, and many cell therapies require product volumes exceeding 100 mL; vitrification has remained largely restricted to small-volume applications because of the technical difficulty of maintaining sufficiently rapid and uniform cooling and warming at larger scales.(4) Thus, previous methods have achieved either clinically relevant processing volumes using slow freezing or rapid, low-CPA vitrification at small scale, but not both simultaneously. The key challenge in vitrification implementation hence resides in maintaining high cooling and warming rates, all while cryopreserving cells in a scalable and continuous manner. A promising route to achieve all of these goals is via sprays, as spray processes increase the surface area-to-volume ratio and offer the potential for high cooling rates. This advantage has historically been limited; small droplets sprayed directly into liquid nitrogen interact disfavorably with the liquid interface due to the inverse Leidenfrost effect and surface tension. Continuous boiling of the liquid nitrogen at the moment of contact keeps small droplets (<30µL) from impinging into the liquid, drastically lowering cooling rates.(37) Moreover, floating or levitating droplets can collide and coalesce at the nitrogen surface, further reducing effective cooling rates and making sub-millimeter droplets impractical for direct spraying into liquid nitrogen. Effective heat transfer rates are larger for droplets exceeding this size threshold, but result in overall lower cooling rates due to the increased thermal capacitance of such droplets.

Previous studies of convection-based droplet printing show droplets on the order of 180 pL (70 µm diameter) achieve estimated cooling rates of approximately 25,000 °C min^−1^, whereas droplets near 520 pL in volume (100 µm diameter) cool at roughly 1,000 °C min^−1^.(26, 27) This reduction in cooling rate was accompanied by a corresponding decline in post-warming viability, from 90% to 79%, which can be attributed to the higher required CPA concentrations assuming a complete absence of ice on cooling and warming. Despite this reduction in viability, increasing droplet size also markedly enhances processing throughput, rising from only 0.5 mL h^−1^ for 70 µm diameter droplets to about 250 mL h^−1^ for 100 µm diameter droplets, illustrating how current gains in scalability come at the expense of cooling efficiency and cell survival. Experimental studies in cell vitrification illustrating this size-dependent trade-off are shown in Figure 1. To circumvent the inverse Leidenfrost effect, several cryopreservation methods rely on spraying or printing droplets directly onto a chilled surface, utilizing conduction cooling to generate rapid cooling rates (10^4^ − 10^6^ ◦C min^−1^).(25, 37, 39, 40) Recovery rates range from 71-95%, with more rapid cooling and warming generally associated with higher viability post-cryopreservation. Although these methods exhibit high viability post-cryopreservation (71-95%), they are severely limited in practical utility because they are batch operations; once the conductive substrate is coated in droplets, the process stops until a new substrate is added. This batch-limited throughput renders them infeasible for the large-scale cryopreservation of next-generation cell-therapy products (typical volumes 10 mL to 100 mL).(4, 41) For example, the surface area of the substrate required to vitrify 1L of the (40 pL) droplets in Akiyama et al. (25) is immense, occupying approximately 100 m^2^. For larger 10µL droplets, the area required is approximately 1 m^2^.

**Fig. 1.**
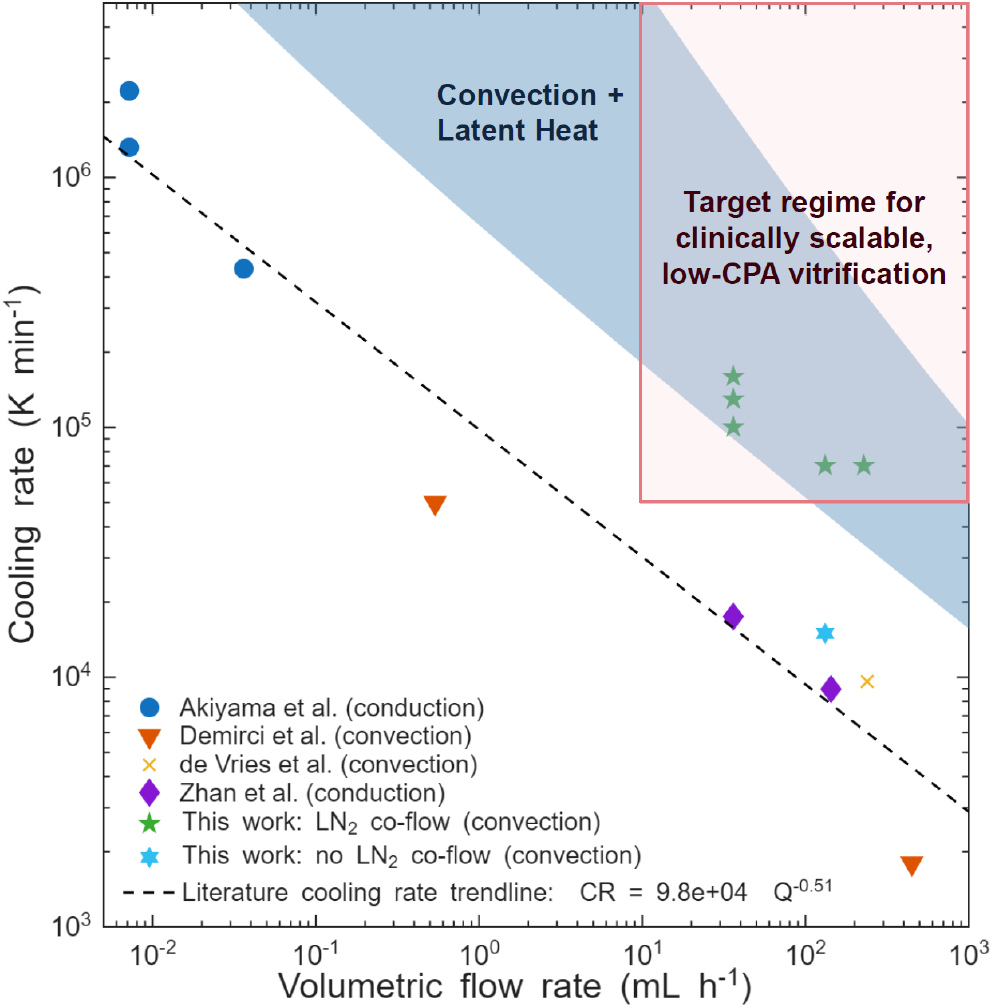
Cooling rate–volumetric throughput map for cryopreservation via droplet vitrification from prior studies, including the work presented here. Cooling rates from literature values were reported from the source material or estimated via analysis of critical cooling rate models for CPAs presented prior.(25–29) Mapping reveals an apparent tradeoff in prior droplet-vitrification approaches between cooling rate and volumetric throughput (dashed line), limiting access to the pink-shaded target regime for clinically scalable, low-CPA vitrification (> 10 mL h^−1^ and > 50,000 K min^−1^); the throughput threshold is motivated by cell-therapy products that can exceed 100 mL and by practical limits on increasing cell concentration to reduce product volume.(4) Also shown are expected cooling rates based on heat transfer modeling of achievable cooling rates in the presented cryoaerosolization process, accounting for both convection and liquid nitrogen latent heat release.

As evidenced in Figure 1, prior droplet vitrification studies define an apparent tradeoff between cooling rate and volumetric throughput. Data from Akiyama et al. (25), Demirci and Montesano (26), de Vries et al. (27), and Zhan et al. (37) collapse onto an approximately power-law trend, with higher throughput generally achieved at the expense of lower cooling rates. In contrast, our cryoaerosolization measurements lie above this trend, achieving substantially higher throughput than prior high-cooling-rate approaches while maintaining cooling rates above those reported for convective droplet vitrification methods. Although the cooling rates remain below the extreme values reported by Akiyama et al. (25), cryoaerosolization operates at much higher volumetric throughput and achieves nearly an order-of-magnitude improvement in cooling rate relative to prior systems with comparable throughput, increasing from approximately 25 000 °C min^−1^ to 210 000 °C min^−1^. Heat-transfer modeling supports this interpretation: gas-side convection alone cannot account for the observed cooling rates, whereas inclusion of latent heat removal from entrained LN_2_ droplet interactions brings predicted rates into the experimentally observed range. This combined convection–latent cooling mechanism enables cryoaerosolization to shift the cooling-rate–throughput tradeoff, maintaining cooling rates in excess of 1 × 10^5^ K min^−1^ while processing droplets continuously at flow rates up to 250 mL h^−1^. To demonstrate biological utility, we applied this platform to vitrify and rewarm human dermal fibroblasts, human induced pluripotent stem cells, and red blood cells at reduced CPA concentrations while maintaining high post-warming viability, with preserved colony-forming ability additionally demonstrated for hiPSCs. To our knowledge, this is the first droplet vitrification-based cryopreservation platform to combine rapid cooling and warming with continuous, clinically relevant throughput, opening the door to high-fidelity cryopreservation of the large cell volumes required for current and future cell therapy products.

## 1. Results and Discussion

### A. Cryoaerosolization

Aerosolization was accomplished using a vibrating orifice aerosol generator (VOAG)(42), which produces highly monodisperse droplets by driving liquid through a small orifice while applying controlled mechanical vibration. CPA with or without cells was supplied to the VOAG nozzle head through syringe tubing and exited through a stainless steel orifice as a continuous microjet (Figure 2a,b). Mechanical oscillations imposed periodic disturbances on the liquid jet, which amplified through the Rayleigh–Plateau instability and caused the jet to break into discrete droplets at regular intervals. The resulting droplet diameter was determined primarily by the orifice size, volumetric flow rate *Q*, and vibration frequency *f*, following the approximate relationship(42):

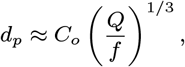

where *C*_*o*_ is a constant that depends on the orifice geometry. The droplet stream was transported downward by a dispersion-air co-flow, which helped maintain droplet separation and prevent aggregation or coalescence. This low-shear aerosolization process is well suited for producing cell-laden droplets for cryopreservation.

**Fig. 2.**
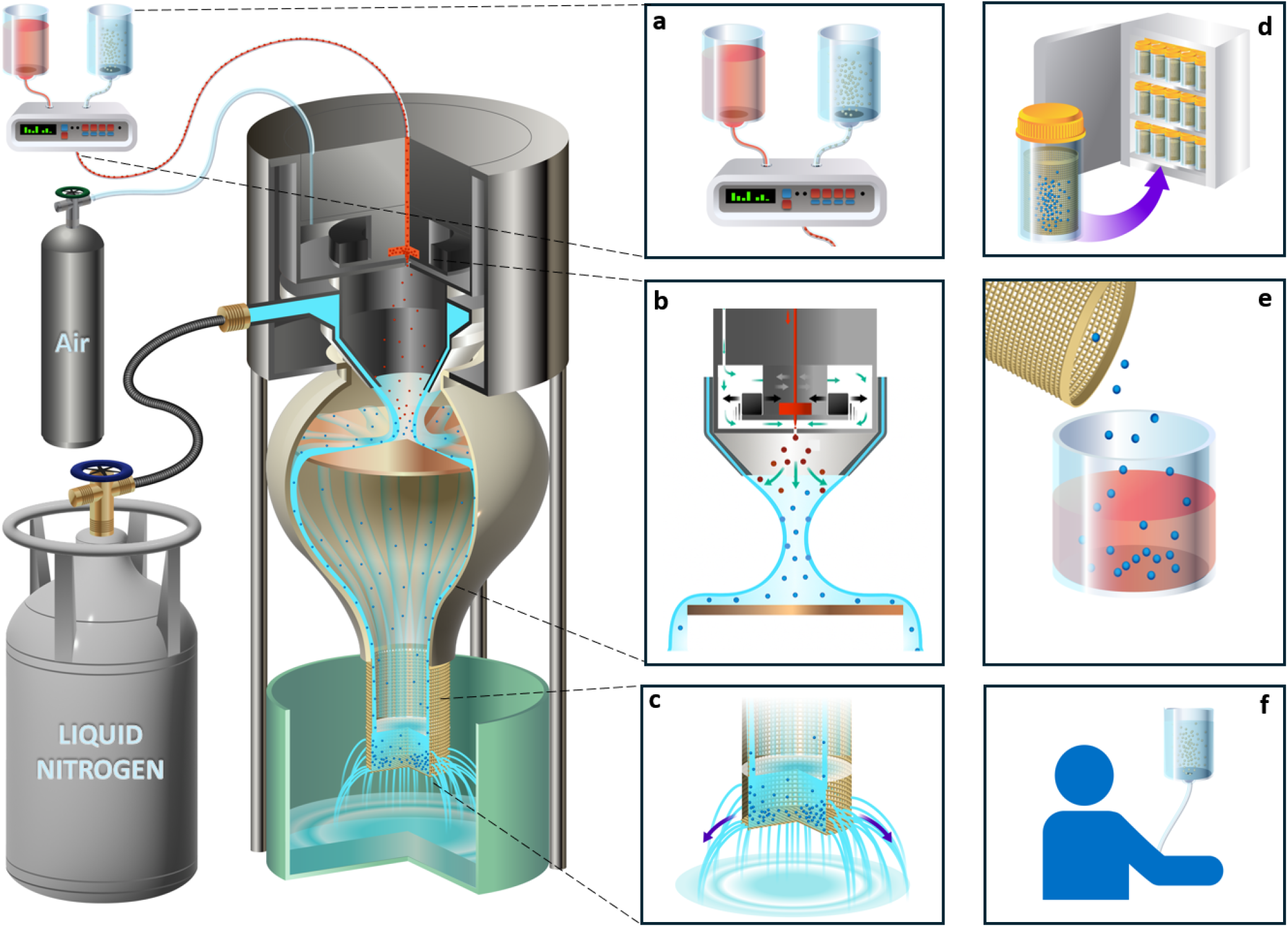
Schematic diagram of the developed high-throughput cryopreservation process. (a) Cells are combined with CPA continuously in-line prior to aerosolization, with variable exposure times controlled by tubing length. The cell-CPA solution then passes through a piezoelectric vibrating orifice (35-200 µm diameter) (b), creating a monodisperse bioaerosol. The bioaerosol then passes through an impinging conical stream of liquid nitrogen (also b), cooling the droplets extremely rapidly (up to 200,000 °C min^−1^), vitrifying the droplets. The droplets are subsequently collected in a 10 µm mesh cylindrical collector (c), which can then be placed in a 50 mL Falcon tube and can contain tens of millions to several billion cells, depending on cell type and initial cell concentration, before transfer to long-term storage (d) or shipping. Rewarming occurs by introducing cell-laden droplets to rewarming solution (e), producing a diluted cell suspension that could then be used for downstream processing or infusion (f). Since the droplet volume fraction is much lower than the rewarming-solution volume (<5%), the final CPA concentration is approximately 1%.

The VOAG was tested with orifice diameters spanning 35–200 *µ*m during development, while the quantitative characterization reported here used 50, 100, and 150 *µ*m orifices. For these characterization experiments, liquid flow rates ranged from 0.6–3.8 mL min^−1^, vibration frequencies from 2.25–17.25 kHz, and dispersion-air flow rates from 1.5– 5.0 L min^−1^; the full operating conditions are provided in Table S1.

Spraying droplets directly into liquid nitrogen can lead to suboptimal cooling because small droplets tend to levitate at the liquid-nitrogen interface rather than immediately penetrating into the liquid bath(37). To increase the cooling rates experienced by aerosolized droplets, we designed an in-flight liquid-nitrogen impingement nozzle (Figure 4b–d). Liquid nitrogen was transferred from a bulk tank to the nozzle through a vacuum-jacketed transfer hose to minimize boiling during delivery. Upon entering the conical nozzle, the liquid nitrogen formed a thin conical spray composed of liquid-nitrogen droplets entrained in cold nitrogen vapor. This spray surrounded the VOAG droplet jet, increasing contact between the aerosolized droplets and the liquid nitrogen. Rapid cooling (>100,000 °C/min) was achieved through a combination of forced convection and droplet–droplet and droplet–film interactions, with additional heat removal from liquid-nitrogen vaporization during collisions.

After passing through the liquid nitrogen spray, the CPA droplet stream impacted a pre-chilled 1.5 mm-thick copper plate, which slowed the droplet trajectories and promoted continuous capture of the solidified droplets. Without the copper impactor plate, droplets tended to adhere to nearby surfaces as they transitioned from supercooled liquid to an amorphous glass. The impactor plate also helped break up larger droplets, which can increase subsequent warming rates because smaller droplets rewarm more rapidly during convective warming. Droplets that were not fully cooled during flight were further cooled by liquid nitrogen on the plate surface before collection. The droplets were then collected in a cylindrical copper mesh tube with a pore size of 10 *µ*m (Figure 2c), allowing liquid nitrogen to drain while retaining the solidified droplets. Previous designs using the liquid-nitrogen spray without the impactor plate achieved similar cooling rates but resulted in greater aggregation and less consistent cooling at higher CPA solution flow rates (Figure 4j). After vitrification, the mesh tube containing the vitrified droplets was placed in a 50 mL centrifuge tube and transferred to liquid-nitrogen storage or deep-cold storage at −150◦C (Figure 2d).

### B. Aerosol Cooling Characterization

Due to the stochastic nature of aerosolized droplet collisions with liquid nitrogen droplets and variability introduced by the inverse Leidenfrost effect, droplets undergoing cryoaerosolization do not experience a single uniform cooling rate, but rather a distribution of cooling rates. This behavior is likely present in many liquid-nitrogen-based cooling systems, although it is often not addressed explicitly in measurement interpretation. Because each experiment involved tens of thousands of droplets, we were able to estimate statistically robust cooling-rate distributions under different operating conditions. To quantify these distributions, we developed a critical cooling rate (CCR) inversion analysis that estimates the underlying cooling-rate probability density from measured crystallized fractions. In this approach, droplets were collected and characterized using a dark-field optical ice assay as described previously(29). Droplets that appeared transparent were classified as vitrified, whereas opaque droplets were classified as crystallized. Previous studies corroborate this method as approximately consistent with measurements made with X-ray diffraction.(28) Controlled illumination and standardized image thresholding ensured consistent characterization across all trials. To handle the large number of images acquired, we implemented a deep neural network-based image-analysis algorithm (Figure 3). The network was trained on manually labeled images containing both vitrified and frozen droplets of varying sizes. After training, it automatically segmented individual droplets, after which droplets were classified using standardized image-intensity thresholds (Figure 3e,f). This approach enabled rapid and reproducible quantification of the crystallized fraction required for the CCR inversion analysis.

**Fig. 3.**
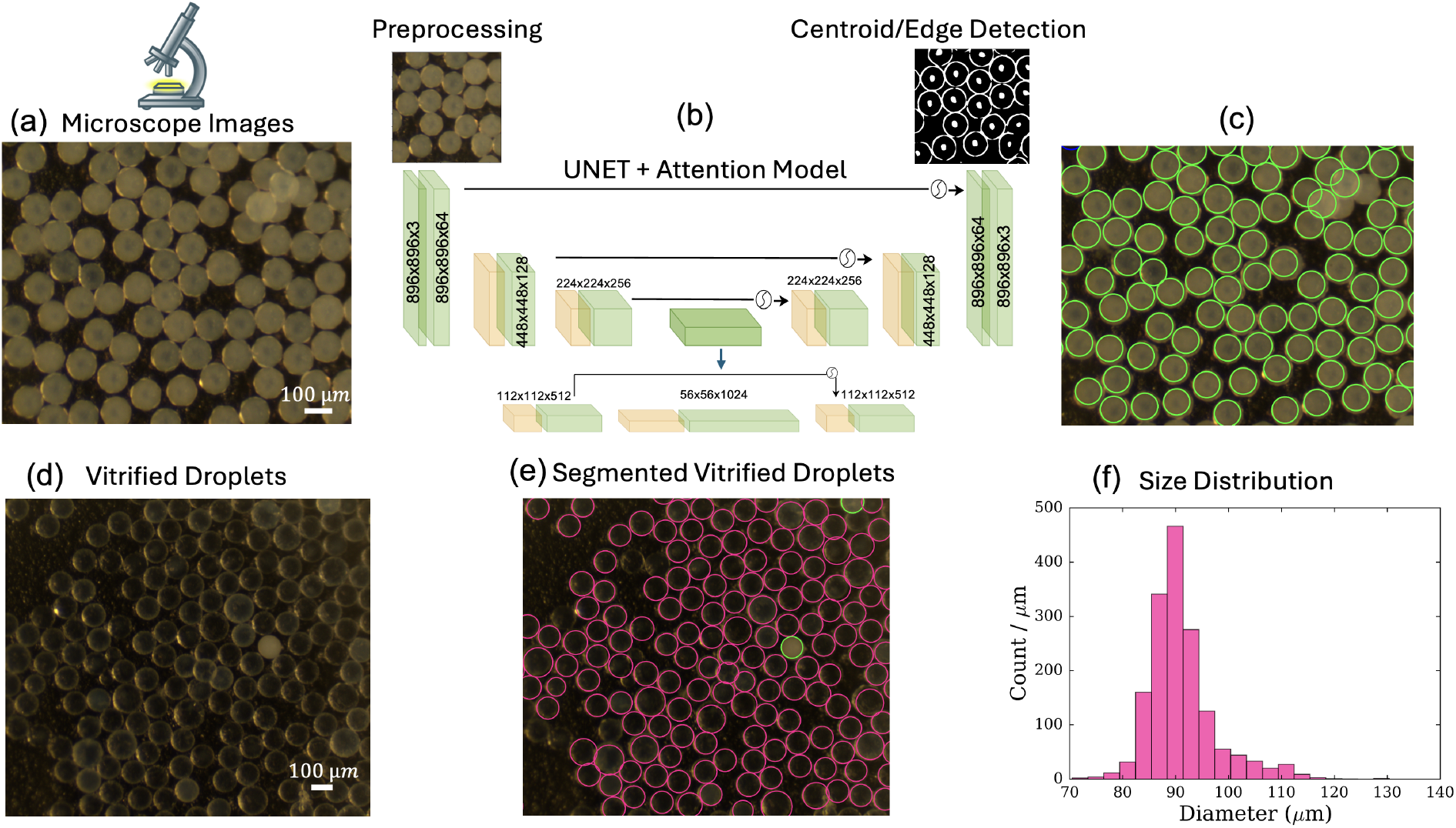
Schematic diagram depicting the image processing pipeline to classify and calculate the diameter of droplets. (a) Sprayed droplets are imaged under a stereo microscope, (b) the images are resized and cropped into fifteen images and then fed to a pretrained deep neural network. The neural network predicts the centroid and edge/perimeter of the droplets, which are further processed to classify droplets and determine droplet diameters. The neural network can handle droplets of different signal intensity, including crystallized (c) and vitrified droplets (d), and segments them for diameter analysis (e). This droplet diameter detection algorithm yields the droplet size distribution (f).

The crystallized fraction, *f* (*c*), was calculated as the droplet volume-weighted fraction of crystallized droplets at concentration *c*, with *f* (*c*) = 0 corresponding to a fully vitrified population and *f* (*c*) = 1 corresponding to a fully crystallized population. For a given cryoprotective solution, the measured dependence of crystallized fraction on concentration was fit using

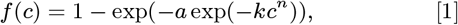

where *a, k*, and *n* are fitting parameters related to the critical cooling rate and crystallization tendency of the droplet solution. Because each concentration corresponds to a specific critical cooling rate, the measured crystallized fraction reflects the fraction of droplets that experienced cooling rates below that threshold. By scanning through CPA concentrations, we obtained samples of the cumulative cooling-rate distribution, which were inverted to reconstruct the underlying cooling rate probability density. This CCR inversion approach enables estimation of cooling-rate distributions directly from binary vitrification outcomes in systems where direct thermal measurements are not possible.

We next evaluated how cooling rate distributions could be modulated during cryoaerosolization. The two primary experimental parameters influencing cooling rate were the liquid nitrogen flow rate through the cooling nozzle and droplet diameter, which was adjusted through orifice diameter and VOAG vibration frequency. To probe the effective cooling-rate distributions, we aerosolized water–ethanol mixtures; ethanol was selected because it is a CPA with crystallization behavior at low concentrations similar to that of other more common CPAs, and because it evaporates from surfaces without leaving residue. Aqueous ethanol solutions ranging from 0.10–0.25 w/w were cryoaerosolized through a 50 *µ*m orifice, corresponding to approximately 90 *µ*m diameter droplets (Figure 3f). Increasing the liquid-nitrogen flow rate increased both the slip velocity between the aerosolized droplets and liquid nitrogen droplets and the convective cooling contribution, resulting in higher cooling rates. Figure 4b–d shows images of the cooling nozzle operated at liquid nitrogen flow rates of 269, 537, and 808 g min^−1^, respectively, with the VOAG droplet jet shown in Figure 4a for reference. Figure 4e shows that increasing liquid nitrogen flow rate increased the mean cooling rate. The mean cooling rate increased from approximately 10,000 K min^−1^ for direct spraying into liquid nitrogen to approximately 210,000 K min^−1^ at the highest liquid nitrogen flow rate. Based on critical cooling rates for propylene glycol solutions, this increase corresponds to a reduction in the CPA concentration required for vitrification from 0.255 w/w to 0.18 w/w.

**Fig. 4.**
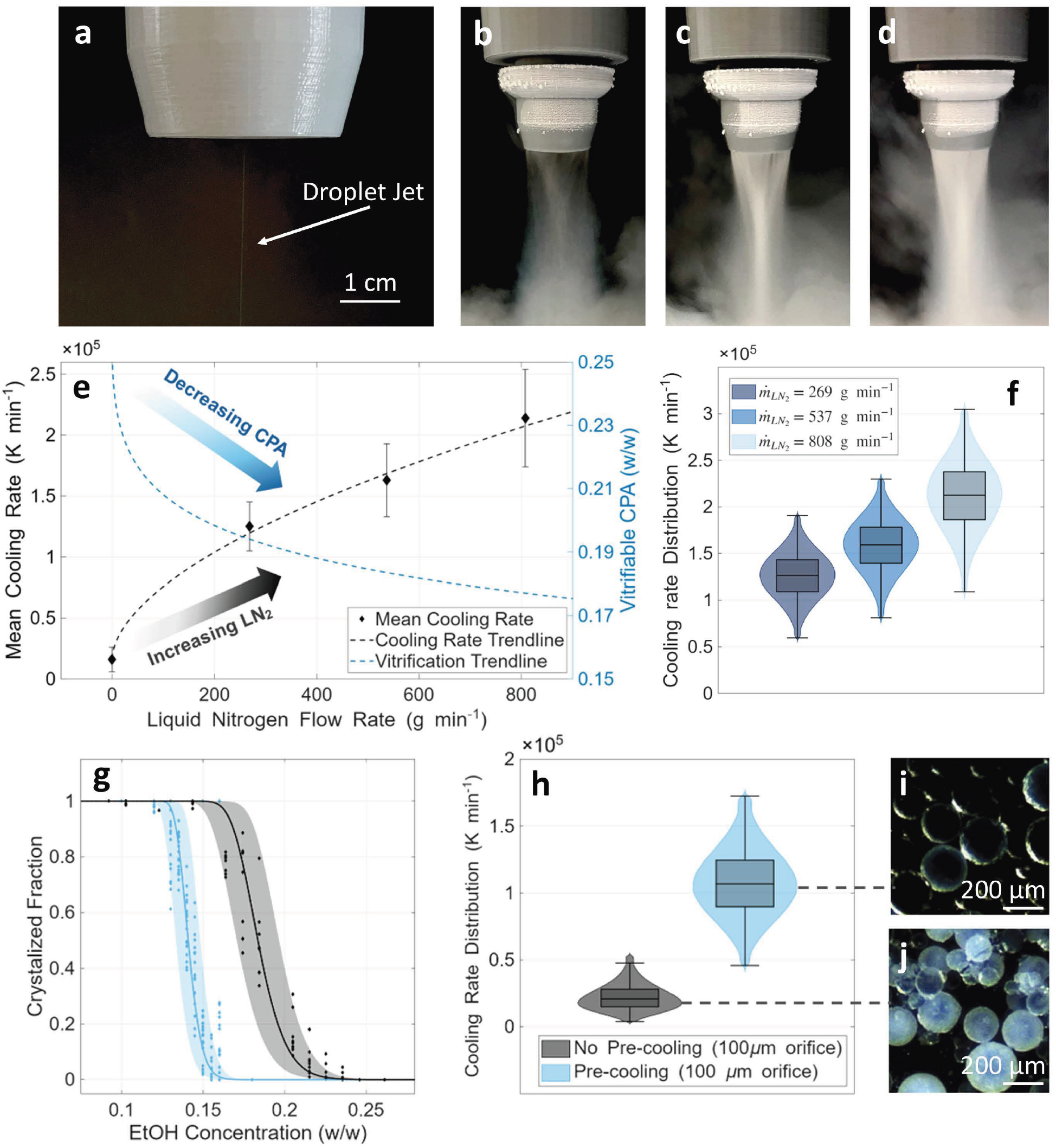
Droplet jet production as it passes through the liquid nitrogen nozzle (a). Images of cryoaerosolization operation at three liquid nitrogen flow rates: (b) 269 gmin ^−1^, (c) 537 gmin ^−1^, (d) 808 gmin ^−1^. (e) Effect of liquid nitrogen flow rate on mean cooling rate and vitrifiable CPA, with the general trend of increasing liquid nitrogen flow rate increasing the mean cooling rate and decreasing the necessary CPA concentration for vitrification. Data are shown for droplets generated with a 50 µm diameter orifice. (f) Cooling rate distributions associated with the three liquid nitrogen flow rates for a 50 µm diameter orifice. (g) Crystallized fraction vs ethanol concentration for droplets produced from cryoaerosolization through a 100 µm diameter orifice. (h) The effect that the pre-cooling nozzle has on cooling rate distribution and the associated (i) vitrified and (j) frozen droplets produced from cryoaerosolization through a 100 µm diameter orifice. The absence of pre-cooling leads to a significant decrease in cooling rates and a more appreciable Leidenfrost effect, causing droplet aggregation on the surface of the liquid nitrogen.

Cooling rate distributions inverted from water–ethanol vitrification data are shown in Figure 4f for liquid nitrogen flow rates of 269, 537, and 808 g min^−1^. Most droplets were estimated to experience cooling rates within approximately ± 20% of the mean, with distribution tails extending to approximately ± 50% of the mean. We attribute the breadth of these distributions to stochastic collisions between aerosolized droplets and liquid nitrogen droplets, as well as spatial variability in slip velocity and local gas flow. Using vacuum-jacketed liquid nitrogen transfer lines helped reduce this variability by lowering the vapor fraction in the nitrogen flow. Future designs may further narrow the cooling rate distribution by improving thermal insulation near the nozzle and incorporating an inline phase separator to purge vapor from the liquid nitrogen feed line.

We also aerosolized aqueous ethanol solutions through a 100 *µ*m orifice, corresponding to approximately 200 *µ*m diameter droplets. Crystallized fraction measurements were obtained for droplets sprayed with and without liquid-nitrogen pre-cooling before entering a liquid nitrogen bath, without the copper impactor plate (Figure 4g). Using the liquid-nitrogen pre-cooling nozzle increased the mean cooling rate from approximately 18,000 K min^−1^ to 110,000 K min^−1^ (Figure 4h). Representative images for the liquid-nitrogen flow case and no-flow case are shown in Figure 4i and Figure 4j, respectively. The liquid nitrogen flow case produced a roughly monodisperse droplet population, whereas the no-flow case produced a broader size distribution. We attribute this difference to droplet levitation at the liquid nitrogen surface through the inverse Leidenfrost effect, which promotes aggregation and reduces effective cooling rates. Aggregation also decreases warming rates during rewarming, further reducing the likelihood that droplets can be vitrified, stored, and rewarmed without crystallization.

### C. Stresses Encountered During Aerosolization

Aerosolization subjects the fluid and encapsulated cells to several distinct stresses (summarized in Figure 5) which must be considered with regard to their influence on cell viability. Inside the orifice, viscous wall shear stress *τ*_*w*_ arises, scaling strongly with orifice diameter *D*. For liquid flow rate *Q* and dynamic viscosity *µ, τ*_*w*_ ∼ *µQ/D*^3^. As the liquid jet is perturbed by the piezoelectric transducer, breakup into droplets occurs through the Rayleigh–Plateau instability, which imposes oscillatory extensional stresses. These extensional stresses scale as *τ*_ext_ ∼ 3*µf*, increasing linearly with vibration frequency *f* and largely independent of jet velocity. For typical VOAG settings, this corresponds to stresses between 100-500 Pa over 10-100 µs. These ultrashort exposure times are unlikely to substantially reduce cell viability, as can be seen in Figure 7c, where human dermal fibroblasts (HDFs) were aerosolized through orifices varying from 35-150 µm in diameter, with associated piezoelectric frequencies of 3-28 kHz. Orifice size had minimal effect on cell survival for low dispersion air (< 3 L min^−1^).

**Fig. 5.**
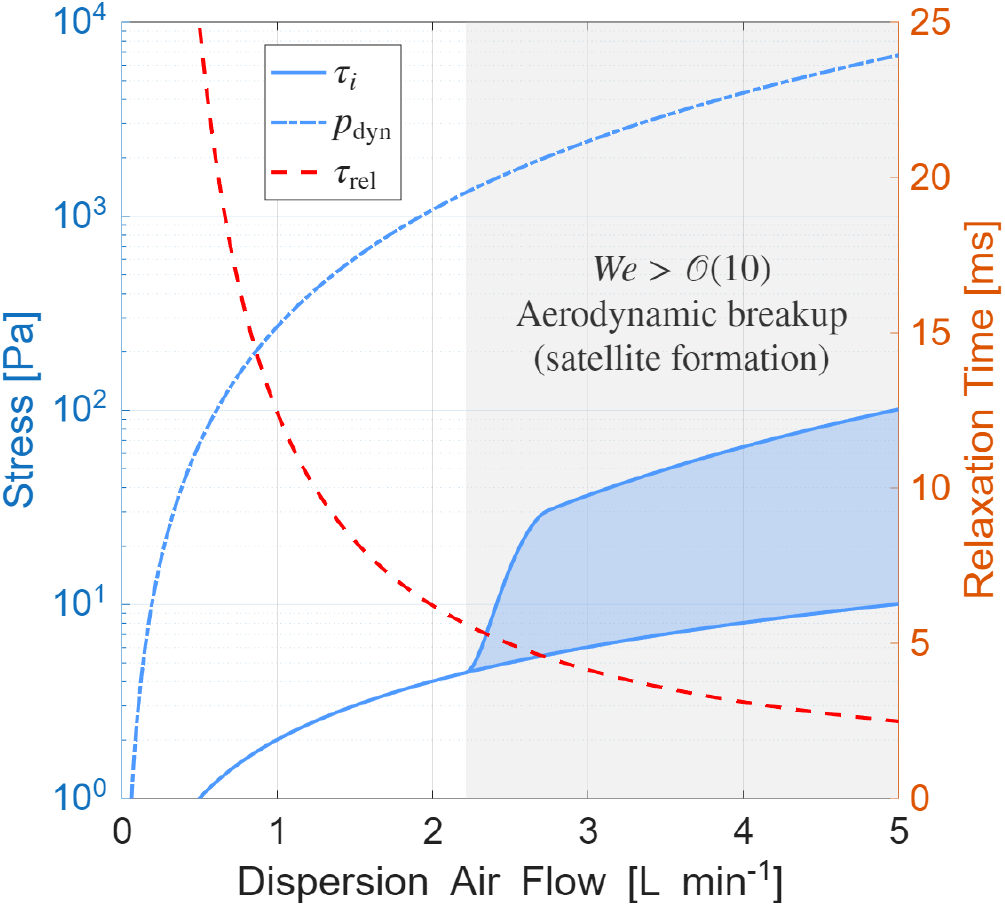
The viscous shear stress *τ*_*i*_, dynamic pressure *p*_dyn_, and relaxation time *τ*_rel_ experienced by the droplets during aerosolization as a function of dispersion air flow rate. The transition between low and high Weber number is also shown, alongside the additional range of viscous shear stresses experienced by the droplets during secondary aerodynamic breakup, which may contribute to the loss in viability at high dispersion air flow rates seen in Figure 7c.

After droplet formation, the dispersion air co-flow introduces an interfacial viscous shear stress, *τ*_*i*_, at the droplet–air interface due to relative motion between the droplet and the surrounding air. This stress has magnitude *τ*_*i*_ ∼ *µ*_*a*_*U*_slip_*/d*, where *d* is the droplet diameter and *U*_slip_ is the *slip velocity*, defined here as the relative velocity between the droplet and the surrounding air. The relevant exposure duration for these air-side stresses is the droplet relaxation time, *τ*_rel_, defined here as the characteristic time required for a droplet exiting the orifice to slow relative to the surrounding gas flow. In parallel, droplets also experience inertial loading from dynamic pressure, 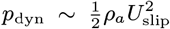. The degree of deformation or breakup is captured by the Weber number, 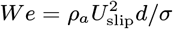, with *We* ≪ 1 indicating nearly spherical droplets and *We* ≳ *O* (10) associated with strong deformation and possible aerodynamic breakup. Dynamic pressure primarily acts as a normal force on the droplet surface and does not directly translate to an intracellular load. As a result, dynamic pressure alone is unlikely to influence cell survival. Its main effect is indirect, by promoting droplet deformation or secondary breakup at sufficiently high Weber numbers, which in turn can expose cells to additional fluid shear during fragmentation. Thus, as shown in Figure 5, air-side stresses become dominant at higher dispersion air flow rates relative to the orifice wall shear and jet breakup stresses described above.

Consistent with our analysis, we observed cell survival was most sensitive to dispersion air flow rate, with larger decreases in viability seen with smaller orifice sizes. We attribute this to decreased droplet size with smaller orifices, reducing the average distance from the cells to the droplet exterior, thereby increasing the effective stresses felt by the cells within the droplet. While the precise stresses are difficult to quantify, the normal operating conditions for cryoaerosolization are well outside the high Weber number regime, with typical dispersion air flow rates around 2 L/min. There may be a benefit to higher dispersion air at larger orifice sizes, as it can reduce droplet size, albeit stochastically, thus increasing cooling and warming rates. Overall, orifice size, applied frequency, and dispersion air all showed minimal effects on HDF viability within the typical operating regime for aerosolization (Figure 7c), indicating the aerosolization process is well tolerated by cells approximately 10µm in diameter. This conclusion is further supported by the minimal viability losses observed for human induced pluripotent stem cells (hiPSCs) and porcine red blood cells (RBCs) after aerosolization (Figure 7a).

### D. Just-In-Time Cryoprotectant Loading

Large-scale cryopreservation of cells and cell-based products requires CPA loading strategies that differ from those used in conventional vitrification techniques. Traditionally, cells are batch-loaded; the entire sample is loaded or step-loaded into CPA prior to vitrification.(37, 43) This approach arises because small sample volumes are vitrified simultaneously, resulting in minimal variation in CPA exposure time between cells, aside from differences introduced by mixing. However, such techniques do not scale to large sample volumes, as processing times may extend to several hours; prolonged CPA exposure during these extended processing periods leads to appreciable cellular damage.(34) Even over a 30-min exposure period, HDF viability decreased by approximately 8% (Figure 6d), with estimated viability decreases of 15% at the one hour mark, with further declines expected at longer times. To circumvent this limitation, we implemented a *just-in-time* CPA loading protocol for HDFs (Figure 6a), in which cells are exposed to full-strength CPA (2.5 M propylene glycol + 1 M trehalose) immediately prior to cryoaerosolization.

**Fig. 6.**
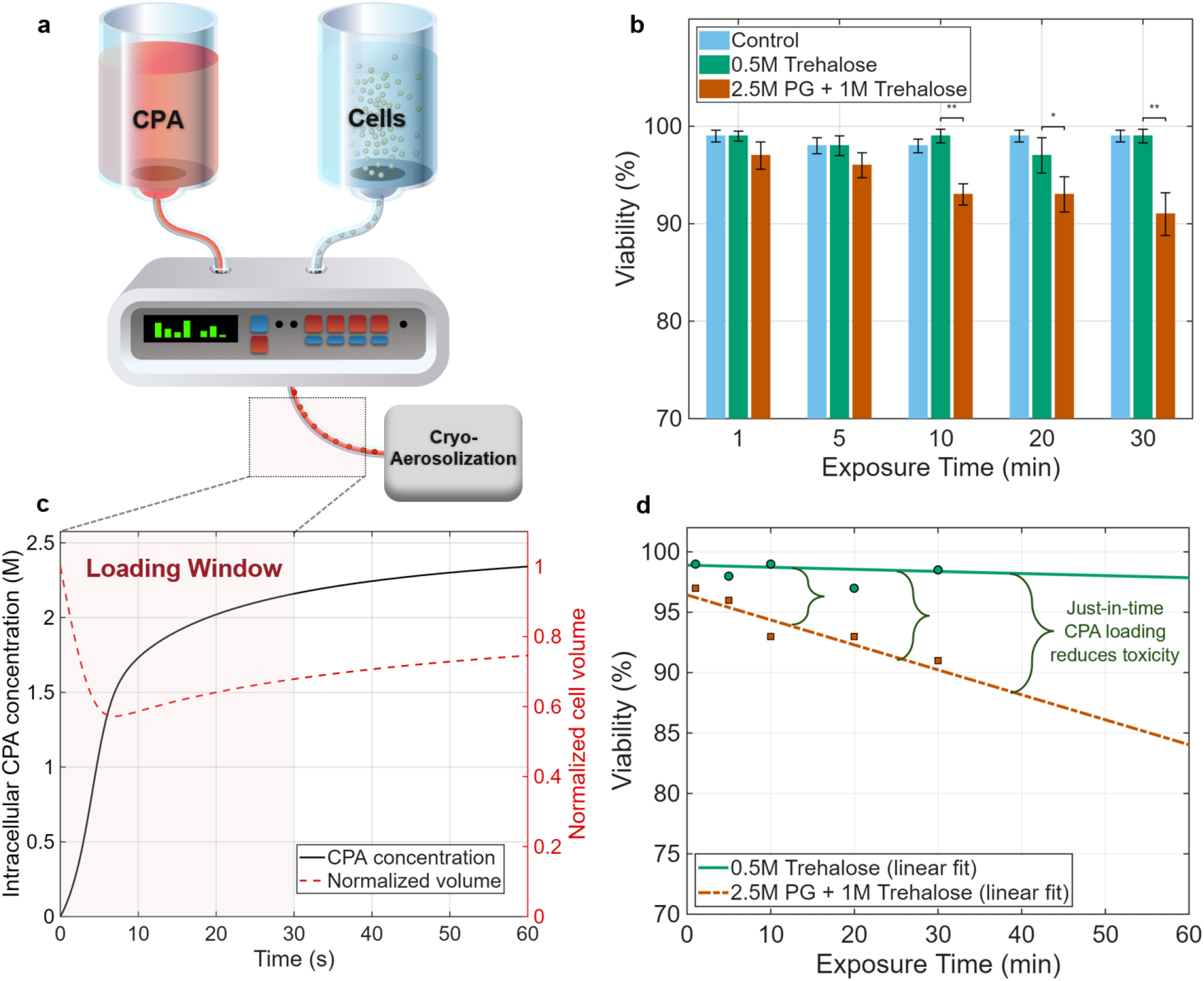
(a) A schematic of the just-in-time CPA loading system, where partially dehydrated HDFs are combined with CPA just prior to cryoaerosolization, allowing for long-term continuous operation without the accrued damage from prolonged CPA exposure. (b) CPA exposure time vs viability for human dermal fibroblasts exposed to the dehydration solution (0.5 M trehalose) and full-strength CPA (2.5 M propylene glycol + 1 M trehalose). (c) Shows normalized HDF volume and internal cell permeating CPA concentration vs time over the 30 s loading window. (d) Shows the difference in HDF viability between prolonged exposure to the full strength CPA vs the dehydration solution, illustrating how a just-in-time loading protocol reduces toxicity over extended periods of time. Statistical analysis was performed using a paired one-way ANOVA followed by Tukey’s post hoc test. Significance symbols: * indicates p < 0.05, and ** indicates p < 0.001.

To reduce osmotic stress during this process, HDFs were pre-loaded with 0.5 M trehalose, a condition that can be maintained for extended periods without appreciable loss in viability (Figure 6b). The dehydrated HDFs and full-concentration CPA were then combined and passed through syringe tubing, with the tubing length selected to limit the transit time, and thus CPA exposure, to approximately 30s, a duration shown by solute transport simulations to be sufficient for near-equilibration of intracellular CPA while minimizing toxic exposure (Figure 6c). Vitrification follows this brief exposure, halting any further CPA-induced toxicity. Figure 6b,d illustrates the advantage of just-in-time CPA loading by showing the dependence of survival on exposure time to both the full-strength CPA and the dehydration solution.

### E. Applications to Cell Cryopreservation

To evaluate the efficacy of this cryopreservation technology across various cell types, we cryoaerosolized and rewarmed hiPSCs, RBCs, and HDFs. Together, these cell types span pluripotent stem cells, blood cells, and adherent somatic cells, representing structurally and functionally distinct populations relevant to regenerative medicine and immunotherapy. Optimization of aerosolization parameters was performed primarily using HDFs and RBCs, which were readily accessible in the quantities necessary for testing stress tolerance. To demonstrate applicability to more sensitive cell types, we evaluated hiPSCs under the optimized cryoaerosolization conditions. These experiments were designed to separate mechanical damage of aerosolization from possible damage during vitrification and rewarming. Cells were aerosolized without cryogenic cooling to determine whether the spray process alone caused appreciable damage. Next, CPA-loaded cells were aerosolized into unloading solution to determine whether stresses from CPA loading amplified those encountered during aerosolization. These same operating conditions were then used for cryoaerosolization and rewarming, allowing the additional effects of vitrification and rewarming to be evaluated directly. This also allowed for comparison with conventional droplet vitrification using matched CPA loading and unloading conditions.

Cryoprotectant formulations and loading strategies were tailored to each cell type to account for differences in membrane permeability and CPA tolerance (see Methods for compositions and protocols). For HDFs, two programmable syringe pumps (SyringePump NE-1000) combined two separate lines immediately prior to aerosolization: one containing full-strength CPA (permeating CPA + 1.5 M trehalose) and the second containing cells at concentrations of 0.5-5 × 10^7^ cells mL^−1^ suspended in 0.5 M trehalose (Figure 6a) maintained at 4◦C. In contrast, RBCs were loaded using a conventional batch protocol at 25◦C due to their ability to tolerate extended CPA exposure without appreciable loss of viability. RBCs were also processed at much higher cell concentrations (approximately 1 × 10^8^ cells mL^−1^) prior to aerosolization. For hiPSCs, conventional step loading and unloading were carried out at 4◦C at an initial cell concentration of approximately 5 × 10^6^ cells mL^−1^.

**Fig. 7.**
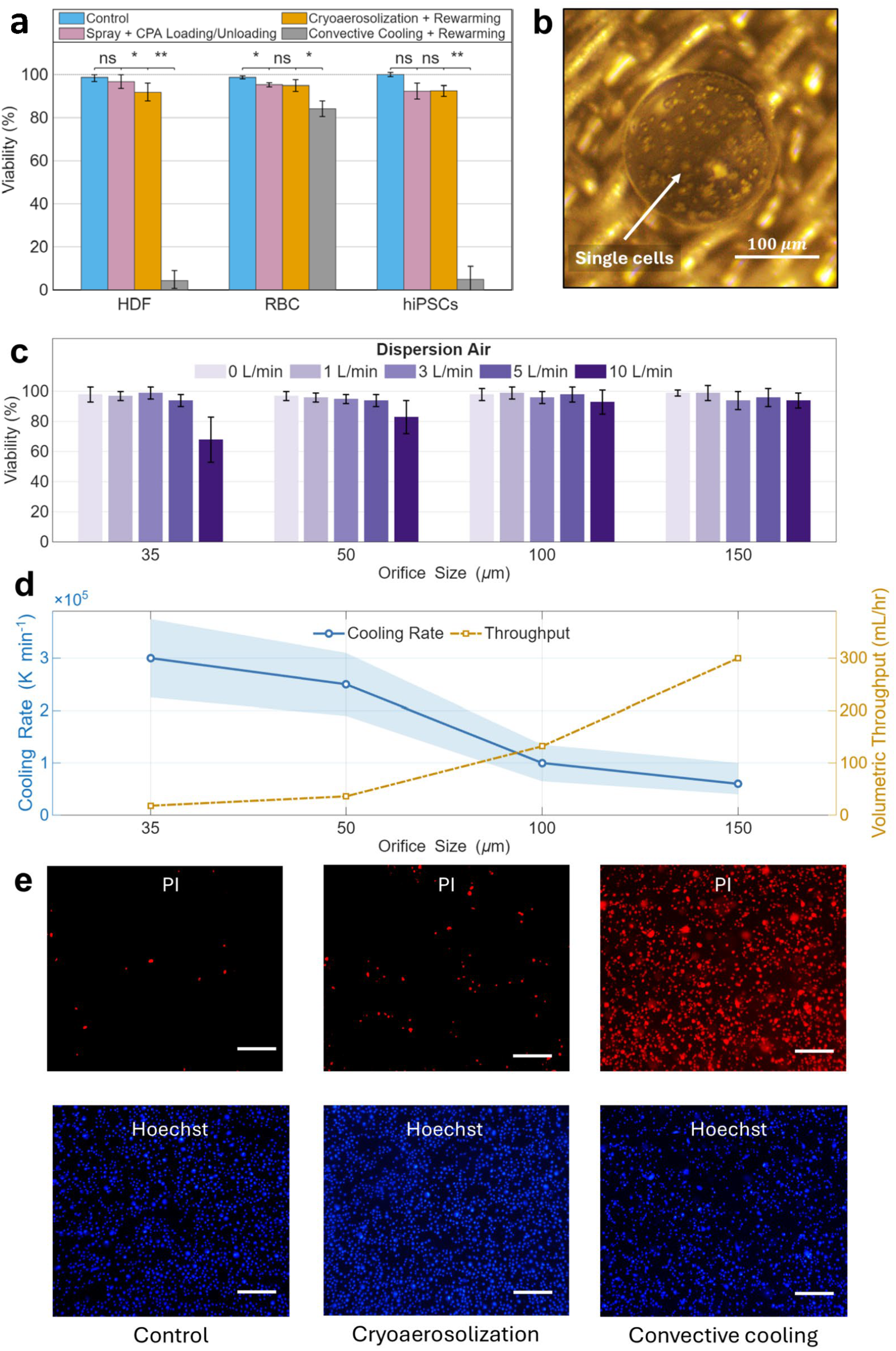
Cryopreservation of hiPSCs (human induced pluripotent stem cells) and HDFs (human dermal fibroblasts) via cryoaerosolization using a 100 µm orifice and for RBCs (red blood cells) using a 150 µm orifice, with cell type-specific cryoprotectant formulations (see Methods). (a) Post-thaw viability/recovery of HDFs, RBCs, and hiPSCs following CPA loading and unloading, cryoaerosolization with rewarming, and conventional droplet vitrification with rewarming (Cryotop). (b) Representative image of a vitrified droplet containing HDFs on the mesh collector following cryoaerosolization. (c) Viability of aerosolized HDFs as a function of dispersion air flow rate and orifice size. The associated piezoelectric frequencies for the 35, 50, 100, and 150 µm orifices were 28 kHz, 17 kHz, 5 kHz, and 3 kHz, respectively. (d) Measured cooling rate and volumetric throughput as a function of orifice size under standard VOAG operating conditions. Data points indicate mean cooling rate; shaded region denotes the 1–99% distribution of measured cooling rates. (e) Representative fluorescence images of Hoechst (all nuclei) and propidium iodide (PI; nonviable cells) for control, cryoaerosolization, and conventional cooling (Cryotop) groups. Scale bars, 200 µm. Data in (a) and (c) are presented as mean ± s.d. Statistical analysis was performed using paired one-way ANOVA with Tukey’s post hoc test. Significance symbols: ns, p > 0.05; *p < 0.05; **p < 0.001.

Following CPA loading, a range of aerosolization parameters was evaluated using HDFs to identify operating conditions that minimized mechanical stress during droplet generation (Figure 7c). Viability remained high (>95%) across moderate dispersion air flow rates and orifice sizes, with decreases observed only at the highest dispersion air condition (10 L min^−1^), particularly for smaller orifices, consistent with predicted stress distributions (Figure 5). A 100 µm orifice (5 kHz oscillation frequency) combined with 2.5 L min^−1^ dispersion air was selected as the optimal operating condition, providing stable droplet formation while maintaining maximal cell viability and throughput. Under these optimized conditions, cells were cryoaerosolized to produce droplets with an average diameter of approximately 190 µm. Aerosolization with CPA loading and unloading resulted in >95% viability across HDFs, hiPSCs, and RBCs (Figure 7a), indicating minimal mechanical damage. Viability (membrane integrity) was assessed via Hoechst–propidium iodide (PI) staining and trypan blue exclusion assay for HDFs and hiPSCs and hemolysis analysis for RBCs. After HDF and hiPSC cryoaerosolization, the collected droplets were imaged to assess vitrification and cell–droplet density. Droplets containing both cell types vitrified successfully. HDF-containing droplets had cell counts ranging from approximately 10 to 100 cells per droplet (Figure 7b), whereas hiPSC-containing droplets had lower cell counts of approximately 1–10 cells per droplet due to lower initial cell concentration. Vitrification was assessed via dark-field optical ice assay, whereby oblique light illuminated the droplets, with only ice scattering light back towards the camera(33). At the maximum HDF loading concentration evaluated, the system achieved an estimated cryopreservation throughput of approximately 1× 10^6^ cells s^−1^. Because throughput scales directly with droplet production frequency and cell loading concentration rather than cell phenotype, substantially higher processing rates were achieved for RBC suspensions. For RBC cryoaerosolization, cell densities of approximately 1 × 10^8^ cells mL^−1^ were achieved, and operation at 5.4 mL min^−1^ resulted in an estimated throughput of approximately 1 × 10^7^ cells s^−1^. Vitrification assessment through dark field ice assay was not possible with droplets with high concentrations of RBCs, as they were too opaque for light to transmit through. Although throughput in this study was limited by cell culture logistics, the system is readily capable of operating with multiple nozzles, enabling substantially greater processing speeds.

Rewarming for all cell types was performed by dispensing vitrified droplets into their respective 37 °C rewarming solutions (Figure 2e). Post-rewarming viability following cryoaerosolization with CPA loading and unloading was 91% for HDFs and 90% for hiPSCs, compared to control viabilities of 99% and 98%, respectively (Figure 7a). RBC recovery following cryoaerosolization with CPA loading and unloading was 94%, comparable to CPA exposure alone and consistent with minimal additional damage during the spray and rewarming processes. To assess post-warming function beyond membrane integrity, hiPSCs were reseeded following rewarming. Cryoaerosolized hiPSCs demonstrated high recovery 24 h after seeding and retained the ability to form colonies by day 5, indicating preservation of proliferative and colony-forming capacity following cryopreservation (Figure 8).

**Fig. 8.**
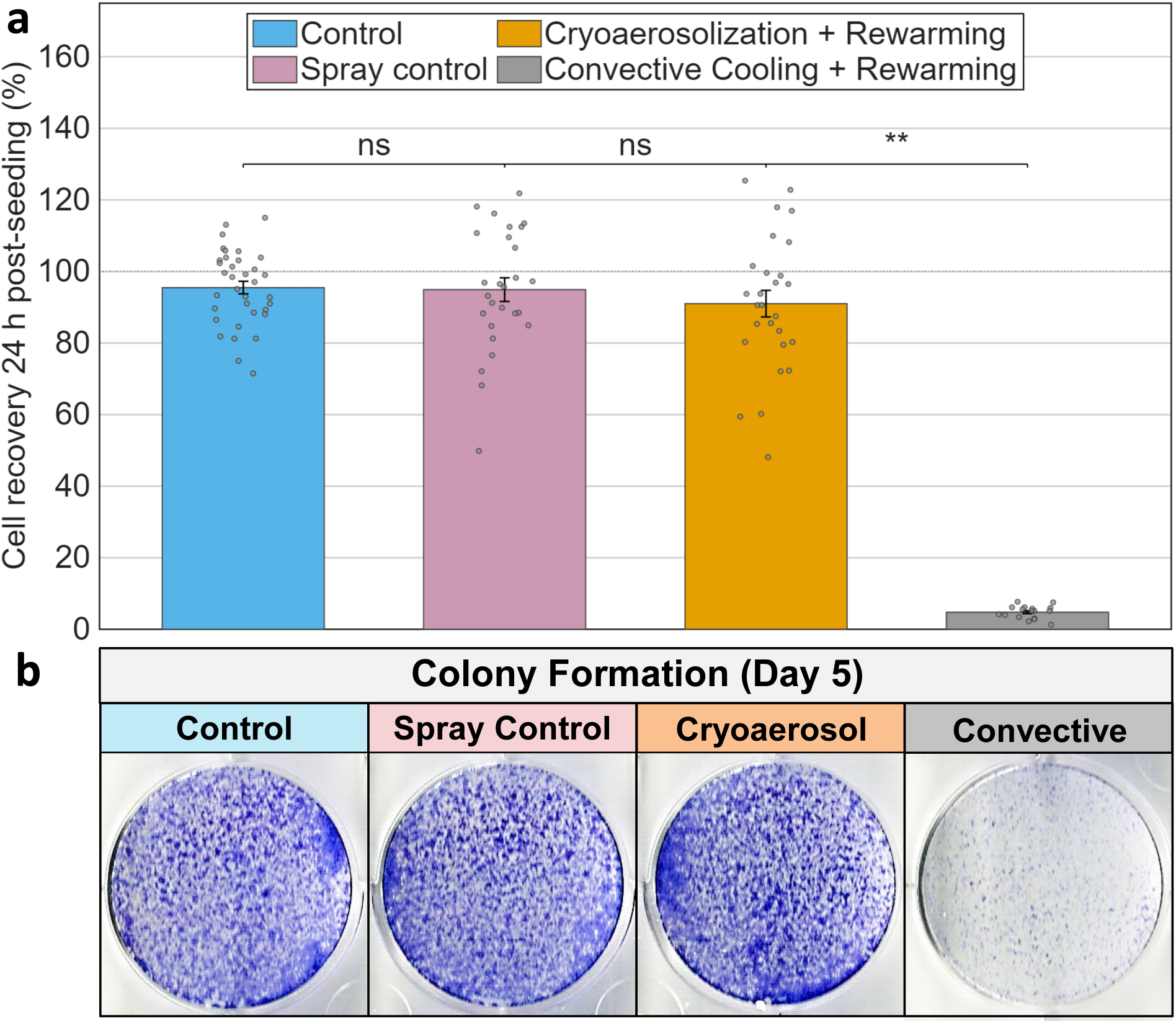
Human induced pluripotent stem cells (hiPSCs) were cryopreserved via cryoaerosolization using a 100 µm orifice, see Figure 7. (a) Shows the cell recovery 24 h post-seeding of 200,000 treated hiPSCs per well. Recovery is given as a percentage of 200,000 for the control group, aerosolization (spray control) only group, cryoaerosolization and rewarming (cryoaerosol) group, and cryotop cooling and rewarming (convective) group. (b) Shows the colony formation at day 5 post-seeding. Data in (a) are presented as mean ± s.d. Statistical analysis was performed using paired one-way ANOVA with Tukey’s post hoc test. Significance symbols: ns, p > 0.05; *p < 0.05; **p < 0.001.

The 50 mL mesh collector provides sufficient capacity for clinically relevant cell numbers. At the cell concentrations evaluated here, a single collector could contain tens of millions to several billion cells, depending on cell type and input concentration. These values are not intrinsic limits of the collector, as the final number of cells stored is determined primarily by the concentration of the initial cell suspension. Following storage, the vitrified droplets can be dispensed directly into a larger volume of rewarming solution, rapidly warming the cells while diluting the CPA. Under the conditions illustrated in Figure 2e, the droplet volume represents less than 5% of the rewarming volume, reducing the final CPA concentration to approximately 1%. For HDFs and hiPSCs, the remaining CPA and extracellular trehalose were subsequently removed by centrifugation and washing before the cells were resuspended in culture medium. Thus, the recovered product is compatible with conventional cell-washing and formulation workflows, although clinical implementation would require these operations to be incorporated into a sterile, closed processing system. For RBCs, glycerol was not removed in the present study, and a standard deglycerolization step would therefore be required before transfusion.

To compare cryoaerosolization with conventional droplet vitrification, each cell type (hiPSCs, RBCs, and HDFs) was also cryopreserved and rewarmed using an in-house fabricated cryotop (consisting of a 100 µm-thick sheet of polypropylene (29)). Cells were first exposed to their respective CPA formulations (see Methods) for 5 min at 4 °C. For hiPSCs and HDFs, 4 µL droplets were placed on the cryotop tip and plunged directly into liquid nitrogen, whereas RBC suspensions were vitrified as 20 µL droplets due to their higher cell loading concentration. Post-cooling assessment using dark-field optical analysis confirmed successful vitrification of the HDF and hiPSC droplets.

Despite successful vitrification, convective rewarming of cryotop cryopreserved samples yielded substantially lower post-thaw viability for nucleated cells, with survival of 4% for HDFs and 5% for hiPSCs, indicating ice formation upon rewarming. In contrast, RBCs exhibited higher post-thaw recovery (84%), consistent with their greater hydraulic conductivity and tolerance to ice formation and the related stresses accompanying it. These results highlight the limitations of conventional droplet vitrification and convective rewarming at microliter scales, particularly for sensitive cell types.

### F. Droplet Warming Simulations

Due to the transient nature of rewarming, as well as the small droplet sizes involved in cryoaerosolization, direct measurement of the warming rates involved is not feasible. Instead, we rely on computational simulations to determine the warming rates involved in our study. Simulations of warming rates (see Experimental) indicate rates in excess of 5×10^5^ K min^−1^ for 200 µm-diameter droplets and 1 × 10^6^ K min^−1^ for 100 µm droplets (Figure 9). Exterior rewarming solution temperatures measured 5 µm from the droplet surface dip slightly below the melting point of 1 M trehalose for very short durations, on the order of 10 ms. Since trehalose itself is a cryoprotectant with a critical cooling rate of approximately 1 × 10^5^ K min^−1^ (at 0.5 M), crystallization of the rewarming solution is not expected to occur or interfere with droplet rewarming. Some uncertainty remains regarding transient thermal behavior prior to droplet contact with the rewarming solution; therefore, the values reported here should be interpreted as estimates of the upper bound on achievable warming rates.

**Fig. 9.**
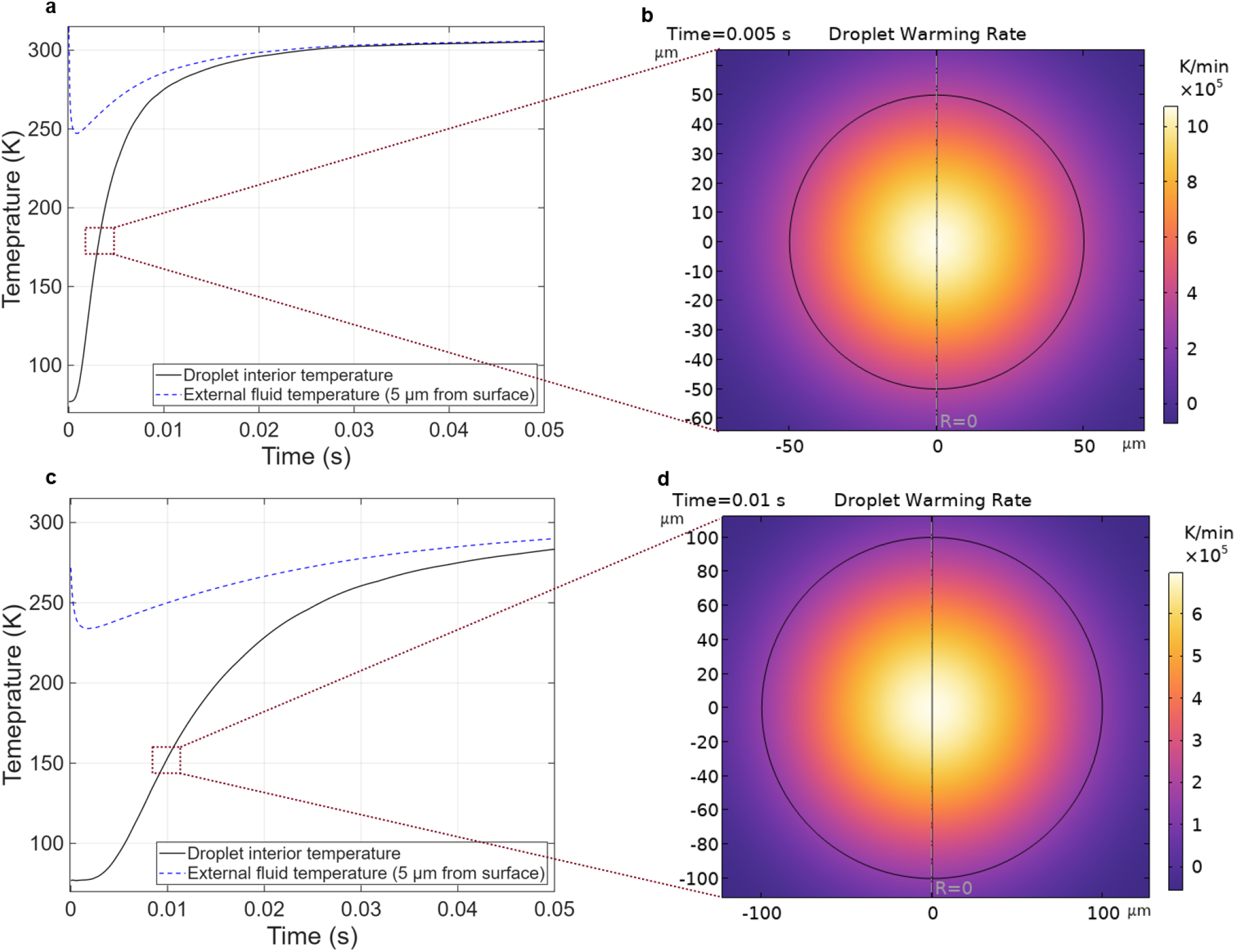
Rewarming simulations of vitrified droplets in contact with a 37 ◦C, 1 M trehalose solution. (a,c) Temporal evolution of the droplet interior temperature (solid line) and the external rewarming solution temperature measured 5 µm from the droplet surface (dashed line) for (a) 100 µm-diameter and (c) 200 µm-diameter droplets. (b,d) Spatially resolved instantaneous warming rates within the droplets at representative times for (b) 100 µm-diameter and (d) 200 µm-diameter droplets. All simulations assume thermophysical properties corresponding to supercooled 2 M propylene glycol.

These rates contrast heavily with those involved for cryotop rewarming. Previous studies indicate rewarming rates for cryotop samples between 1 − 20 µL to be 1− 10 × 10^3^ K min^−1^(29), approximately 2-3 orders of magnitude slower than rewarming rates encountered here. Faster rates nearing 1 × 10^6^ K min^−1^ have been reported for single droplet rewarming (33, 44), though these methods lack the scalability achieved here for broader application in cell cryopreservation.

## 2. Conclusion

This work introduces a new high-throughput cryopreservation technique, cryoaerosolization, that addresses a long-standing issue of achieving rapid cooling and warming rates and clinically relevant throughput. By coupling a vibrating-orifice aerosol generator with an impinging liquid-nitrogen nozzle, micrometer-scale droplets are confined and quenched in flight, reducing cooling losses from the inverse Leidenfrost effect, achieving mean cooling rates up to ∼ 2.1 × 10^5^ ◦C min^−1^ (with ≈ 1 × 10^6^ ◦C min^−1^ on subsequent warming possible) while sustaining processing rates ≳ 100 mL h^−1^. Using cell type-specific CPA formulations containing only 19–25 wt% permeating CPA, HDFs and hiPSCs were cryoaerosolized and rewarmed with approximately 90% post-thaw viability, while RBCs showed 94% post-thaw recovery, all outperforming conventional droplet vitrification operated at matched CPA loading and unloading conditions but lower cooling and warming rates.

Practically, the platform is modular and scalable (multi-nozzle arrays), compatible with *just-in-time* CPA loading to minimize toxicity exposure, and supports straightforward collection, storage, and rapid convective rewarming. These features map directly onto the manufacturing needs of next-generation cell therapies, where large doses, repeat administrations, and logistics favor continuous, closed, and high-rate processes.

Residual variability in cooling arises from stochastic LN_2_–droplet collisions and spatial variability in the flow profile; nozzle-side gas management and phase separation are expected to further narrow cooling rate distributions. While HDFs, hiPSCs, and RBCs provide useful initial test cases, broader validation across sensitive primary cells and engineered lymphocytes, as well as aggregates (e.g., islets, organoids), is warranted. Future work will integrate sterile, closed-loop fluid paths with automated post-rewarming CPA removal and cell concentration, providing a practical route to recover cells at the concentrations required for administration. Closed, automated spinning-membrane systems have already been demonstrated for the harvest, washing, and concentration of activated T cells, tumor-infiltrating lymphocytes, and mesenchymal stromal cells, with satisfactory cell recovery and viability across multiple cell-processing facilities.(45) Multi-orifice arrays will also be implemented to enable liter-scale hourly throughput.

## 3. Experimental

### Cryopreservation protocols

Cell type-specific CPA loading, cryoaerosolization, and unloading protocols were used for HDFs, hiPSCs, and RBCs. For HDFs and hiPSCs, cryoaerosolization was performed using a 100 µm orifice, a liquid flow rate of 2.5 mL min^−1^, a vibration frequency of 5 kHz, a dispersion air flow rate of 2.5 L min^−1^, and a liquid nitrogen flow rate of approximately 500 g min^−1^ through the impinging conical nozzle. RBCs were processed using a 150 µm orifice, a liquid flow rate of 5.4 mL min^−1^, a vibration frequency of 2700 Hz, a dispersion air flow rate of 3 L min^−1^, and a liquid nitrogen flow rate of approximately 500 g min^−1^. Vitrified droplets were collected using an in-house fabricated stainless steel mesh collector with a 50 mL collection volume and 10 µm pore size, then stored in liquid nitrogen prior to rewarming.

For HDF cryopreservation, HDFs at concentrations of 0.5–5 × 10^7^ cells mL^−1^ were loaded using the just-in-time CPA loading protocol. Two programmable syringe pumps (SyringePump NE-1000) were used to deliver separate streams to a T-junction immediately upstream of the aerosolization nozzle. One stream contained CPA solution (5 M propylene glycol + 1.5 M trehalose), while the second contained suspended HDFs in 0.5 M trehalose, maintained at 4 °C using an ice-water bath. The two streams were combined at equal flow rates, yielding a final CPA concentration of 2.5 M propylene glycol and 1 M trehalose immediately prior to aerosolization. The residence time between the mixing junction and the nozzle was controlled by varying tubing length, with a nominal exposure time of 30 s prior to aerosolization. HDFs were then cryoaerosolized under the HDF/hiPSC operating conditions described above, collected in liquid nitrogen, and rewarmed by direct addition to a circulating 37 °C bath containing 1 M trehalose in PBS. Trehalose was removed by centrifugation and washing prior to viability assessment.

For hiPSC cryopreservation, cells were centrifuged and resuspended in 1 mL of phosphate-buffered saline (PBS) to achieve a cell concentration of approximately 5 × 10^6^ cells mL^−1^. The cell suspension was placed on ice, and 14 mL of pre-chilled CPA solution containing 2 M ethylene glycol (EG) and 2 M DMSO in PBS was added dropwise over 3 min, yielding a final concentration of approximately 1.85 M EG and 1.85 M DMSO. Due to the limited supply of hiPSCs, standard CPA loading was used rather than just-in-time CPA loading. CPA-loaded hiPSCs were then cryoaerosolized under the HDF/hiPSC operating conditions described above. Rewarming was carried out in a circulating 37 °C bath containing 1 M EG, 1 M DMSO, and 0.5 M trehalose. Cells were then centrifuged and resuspended in 1 M trehalose for 5 min, followed by a final centrifugation wash and resuspension in hiPSC culture medium prior to subsequent culture and assessment.

RBCs were loaded with glycerol using a stepwise dropwise addition protocol at 20 °C, adapted from Samot et al.(46). RBC suspensions with cell concentration of approximately 4×10^8^ cells mL^−1^ were first mixed 1:1 (v/v) with 2 M glycerol to obtain a final glycerol concentration of 1 M. The partially loaded RBC suspension was then mixed 1:1 (v/v) with 4 M glycerol, yielding a final glycerol concentration of 2.5 M. CPA-loaded RBCs were cryoaerosolized under the RBC operating conditions described above. Vitrified RBC droplets were rewarmed in a circulating 37 °C bath containing 2.5 M glycerol in PBS, matching the final glycerol concentration used during loading. Hemolysis was assessed after each processing step, including CPA loading, cryoaerosolization and rewarming, and conventional droplet vitrification and rewarming. As in Samot et al., glycerol was not removed prior to hemolysis assessment.

### Convective droplet vitrification comparison

For comparison with conventional droplet vitrification and rewarming, CPA-loaded cell suspensions were vitrified by direct plunging on an in-house fabricated Cryotop consisting of a 100 µm-thick polypropylene sheet(29). For HDFs and hiPSCs, 4 µL droplets of loaded cell suspension were placed on the Cryotop and plunged directly into liquid nitrogen. For RBCs, 20 µL droplets were used to match the larger droplet volumes used in the RBC vitrification study reported by Samot et al.(46). After cooling, Cryotop samples were rewarmed in their respective circulating 37 °C warming baths and subjected to the same unloading or washing procedures used for the corresponding cryoaerosolized samples.

### Cell culture and post-warming assessment

Post-warming viability was assessed by trypan blue exclusion and Hoechst– propidium iodide staining for HDFs and hiPSCs, with hiPSC recovery and colony-forming capacity additionally evaluated after reseeding; RBC recovery was assessed by spectrophotometric hemolysis. Human dermal fibroblasts (HDFs; ATCC, PCS-201-012) were cultured in Dulbecco’s Modified Eagle Medium (DMEM) supplemented with 10% fetal bovine serum (Thermo Fisher Scientific) and 1% penicillin–streptomycin (Sigma). Cells were maintained at 37 °C in a humidified incubator with 5% CO_2_ and passaged using 0.05% trypsin– EDTA when cultures reached greater than 85% confluence. Cell viability was assessed using trypan blue exclusion.

Cell suspensions were mixed 1:1 with 0.4% trypan blue solution and quantified using a Countess II automated cell counter (Thermo Fisher Scientific). Viable cells were defined as unstained cells, and viability was calculated as

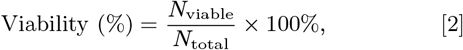

where *N*_viable_ is the number of viable cells and *N*_total_ is the total number of cells counted.

For fluorescence-based live/dead staining, cells were incubated with Hoechst 33342 (Thermo Fisher Scientific, REF#62249) and propidium iodide (PI; Invitrogen, REF#P3566) for 20 min at 37 °C in the dark. After staining, cells were washed once with PBS and imaged immediately using an Olympus IX50 fluorescence microscope at 4 magnification. Images were analyzed using Fiji/ImageJ (version 1.52a). Hoechst-positive/PI-negative nuclei were counted as viable, whereas Hoechst-positive/PI-positive nuclei were counted as non-viable.

Human induced pluripotent stem cells (hiPSCs) were maintained under feeder-free conditions in mTeSR1 medium (STEMCELL Technologies, Cat. #85850) according to the manufacturer’s instructions and to our previous methods.(47) Culture plates were coated with Matrigel (Corning, Cat. #354277) to support cell attachment and maintenance of pluripotency. Matrigel was diluted in cold DMEM/F12 (STEMCELL Technologies, Cat. #36254) according to the manufacturer’s instructions, and plates were incubated at 37 °C for 45 min before use. Excess coating solution was aspirated immediately before cell seeding. hiPSCs were maintained at 37 °C in a humidified incubator with 5% CO_2_, and medium was changed daily.

For routine passaging, hiPSCs were dissociated using ReLeSR (STEMCELL Technologies, Cat. #100-0483) at 60– 70% confluence and gently resuspended to generate small cell aggregates while preserving colony integrity. The resulting cell clusters were replated onto Matrigel-coated plates in fresh complete medium. To generate single-cell suspensions for cryoaerosolization experiments, hiPSCs were dissociated using Accutase (Sigma-Aldrich, A6964). Additional details regarding hiPSC live/dead staining and colony-formation assays are provided in the Supplementary Material, Sections 6 and 7.

To quantify hemolysis, samples were separated into two groups: an experimental group containing RBCs subjected to the experimental condition, and a total-lysis group in which RBCs were mixed with DI water to release the full hemoglobin content. Hemolysis was calculated from the absorbance of the cell-free supernatant as

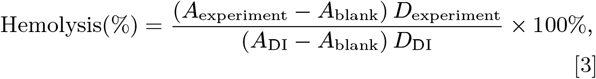

where *A*_experiment_ is the absorbance measured from the experimental sample, *A*_DI_ is the absorbance measured from the total-lysis control, *A*_blank_ is the absorbance of the corresponding blank solution, and *D*_experiment_ and *D*_DI_ are the dilution factors for the experimental and total-lysis samples, respectively. If both samples were diluted identically, the dilution factors cancel.

RBC hemolysis values are shown in Figure 7a. Control RBCs exhibited 98% recovery, where recovery was defined as 100 − hemolysis (%). Following glycerol loading, RBC recovery remained high at 95%, indicating minimal damage from the stepwise loading protocol. After cryoaerosolization and rewarming, RBC recovery was 94%, demonstrating that the cryoaerosolization process introduced little additional hemolysis under these conditions.

### System operation

The system was operated by first setting the desired orifice frequency and syringe pump flow rate to produce the target droplet size for a given orifice. Liquid nitrogen flow was then initiated and adjusted until the target flow rate was reached. The liquid nitrogen flow rate measurement procedure was as follows: (1) the liquid nitrogen flow was initiated and allowed to stabilize for approximately 30 seconds; (2) liquid nitrogen was collected in a pre-weighed, pre-cooled container positioned directly beneath the nozzle outlet; (3) the collection time was measured using a stopwatch, typically for 60 seconds; (4) the container was re-weighed to determine the mass of collected liquid nitrogen; and (5) the mass flow rate was calculated as the collected mass divided by the collection time. If the measured flow rate did not match the target value, the flow control valve was adjusted and the measurement was repeated. To ensure measurement reliability, the procedure was repeated three times for each flow rate setting, and the average value was reported. We also attempted to use a commercial liquid nitrogen flow meter; however, the direct collection method provided more reliable measurements due to the two-phase nature of the flow. The flow meter consistently gave lower readings than the weighing method, likely because a portion of the LN_2_ vaporized after passing through the flow meter.

### Image processing

Droplets were imaged using a stereo microscope under dark-field and bright-field illumination, and full-resolution images of size 5660 × 3680 × 3 pixels were captured, as shown in Figure 3a. Each image was split into fifteen tiles of size 1000 × 1000 × 3. The tiles were resized to 896 × 896 × 3 before input to the pretrained deep neural network to reduce memory usage. During training, an 80/20 train/validation split was used, and a separate held-out test set was maintained for final evaluation.

We used an attention-gated U-Net (48–51) with an encoder–decoder design and skip connections. The encoder contained four blocks; each block applied two 3 × 3 convolutions with ReLU activation and same padding, followed by 2 × 2 max pooling that halved the spatial size. The number of feature maps doubled with depth (64, 128, 256, 512), and a bottleneck operated at 1*/*16 resolution with 1024 channels. For an input tile of size 896 × 896, the encoder resolution progressed as 896, 448, 224, 112, and 56 pixels, while the number of channels increased from 64 to 1024 at the bottleneck. The decoder restored resolution in four steps. At each step, the feature map was upsampled by a factor of two, an attention gate was computed on the corresponding encoder feature map, and the upsampled map was concatenated with the attention-filtered skip connection. A convolutional block then refined the representation and reduced the channel number symmetrically (512, 256, 128, 64) as the spatial resolution returned to 896 × 896. Two parallel 1 × 1 convolution heads with sigmoid activations produced dense probability maps for the droplet edge and droplet centroid. The network was trained end-to-end using the Adam optimizer and binary cross-entropy loss for each head. Dropout with a rate of 0.10 was applied within the convolutional blocks, and batch normalization was used to stabilize training and reduce overfitting. The model contained 37.3 million trainable parameters and was trained using 60 images containing more than 10,000 droplets. The trained model is available in the GitHub repository (52).

Centroid pixels were converted to single droplet centers using the CUDA-accelerated density-based spatial clustering algorithm DBSCAN (53) implemented in cuML. The predicted centroid points were superimposed on the predicted edge channel. From each centroid, the nearest edge was identified by searching in eight directions, and the resulting distances were averaged to estimate the droplet radius. The mean intensity *I* was also measured at each droplet center, and droplets were classified as frozen for *I* > 60 or vitrified for *I* < 60.

We computed the crystallized fraction, *f*_*c*_, following Eq. 4, and fit the data to a sigmoid function, as shown in Figure 4g:

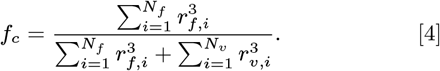

Here, *N*_*f*_ and *N*_*v*_ are the numbers of frozen and vitrified droplets, respectively, and *r*_*f,i*_ and *r*_*v,i*_ are the corresponding droplet radii. Because droplet volume scales with *r*^3^, this expression gives the volume-weighted frozen fraction.

### Cooling-rate distribution

For a given CPA concentration *c*, there exists a well-defined critical cooling rate CCR(*c*) above which droplets vitrify and below which crystallization occurs. When small droplets experience a distribution of cooling rates, the outcome for any individual droplet is therefore approximately binary (vitrified or crystallized), leading to a measurable crystallized droplet fraction that depends on *c*.

Let *R* denote the stochastic cooling rate experienced by a droplet, with probability density *p*_*R*_(*r*). The crystallized fraction measured at concentration *c* is then the probability that the cooling rate falls below the critical value,

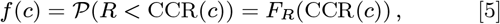

where *F*_*R*_(*r*) is the cumulative distribution function of the cooling rate. Equivalently, the vitrified fraction corresponds to the survival function *S*_*R*_(*r*) = 1 − *F*_*R*_(*r*).

By sweeping the CPA concentration, the critical cooling rate CCR(*c*) is varied, and the measured crystallized fraction directly samples the cumulative distribution of cooling rates at different thresholds. Differentiating with respect to concentration yields

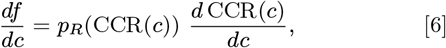

allowing the underlying cooling-rate probability density to be recovered after mapping concentration to critical cooling rate.

Thus, variation of CPA concentration enables inversion of the measured crystallized fractions, allowing for the reconstruction of the cooling-rate distribution experienced by the droplets. Combining the exponential dependence of critical cooling rate with concentration and Avrami kinetics, we arrive at an approximate functional form for *f* (*c*), shown in Eq. 1(22).

### Liquid nitrogen impingement heat-transfer model

Liquid nitrogen exits the conical nozzle as a thin, rapidly boiling sheet into a warmer gas environment. Under these conditions, surface-tension instabilities, aerodynamic shear, and vaporization likely destabilize the film, producing a dense spray of liquid nitrogen droplets rather than a continuous liquid surface. To estimate the cooling rates of droplets as they pass through this dense liquid-nitrogen droplet region, we developed a lumped heat-transfer model for a spherical droplet exposed to cold nitrogen vapor containing entrained liquid nitrogen droplets. The droplet was assumed to have a spatially uniform temperature, such that heat removal occurs through gas-side convection and latent heat absorption during collisions with liquid nitrogen droplets. The transient energy balance is

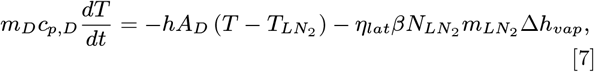

where *m*_*D*_, *c*_*p,D*_, *A*_*D*_, and *T* are the mass, specific heat, surface area, and temperature of the droplet, respectively. The surrounding nitrogen phase was approximated as being at the liquid nitrogen boiling point, 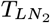 . The first term on the right-hand side represents convective heat transfer, with *h* denoting the gas-side convective heat-transfer coefficient. The second term represents heat removal associated with liquid nitrogen droplet impacts and vaporization, where 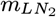 is the mass of an individual liquid nitrogen droplet, Δ*h*_*vap*_ is the latent heat of vaporization of liquid nitrogen, 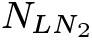 is the liquid nitrogen droplet number concentration, *β* is the collision kernel between liquid nitrogen droplets and aerosolized droplets (54), and *η*_*lat*_ is an effective latent-transfer efficiency that accounts for incomplete vaporization and imperfect thermal coupling during droplet impacts. In the absence of liquid nitrogen droplet impacts, *β* = 0, Eq. 7 reduces to the standard lumped-capacitance convection model (55).

The model was nondimensionalized using

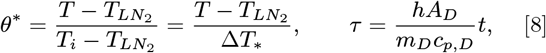

where *T*_*i*_ is the initial droplet temperature and 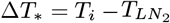. This gives

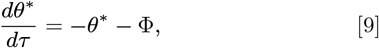

with

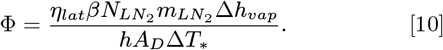

The dimensionless parameter Φ represents the ratio of latent heat removal by liquid nitrogen droplet impacts to gas-side convective heat removal. Thus, Φ = 0 corresponds to convection-only cooling, whereas increasing Φ increases the effective heat-removal rate above that predicted by convection alone. Parameter values used for the calculations shown in Figure 1 are provided in Table S2.

The convection-only model underpredicted the cooling rates inferred from cryoaerosolization experiments, whereas the combined convective–latent model captured the experimentally observed range. This suggests that collisions between liquid nitrogen droplets and aerosolized droplets contribute substantially to heat removal during cryoaerosolization. This additional latent heat-transfer pathway provides a mechanistic explanation for why the present system achieves higher cooling rates at high volumetric throughput than would be expected from gas-side convection alone.

### Solute transport model

Transient cell shrink-swell behavior and intracellular cryoprotectant uptake during just-in-time loading were modeled using a two-parameter membrane transport formalism based on water permeability *L*_*p*_ and solute permeability *P*_*s*_ (56). The total cell volume was written as *V*_*c*_ = *V*_*w*_ + *V*_*s*_ + *V*_*b*_, where *V*_*w*_ is the intracellular water volume, *V*_*s*_ is the intracellular permeating solute volume, and *V*_*b*_ denotes the osmotically inactive volume. Water transport across the membrane was governed by the osmotic pressure difference between intra- and extracellular compartments,

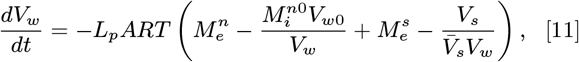

while permeating solute transport was described by simple diffusive permeation,

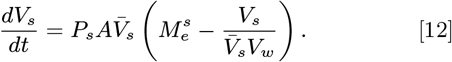

Here *A* is the membrane surface area, *R* is the gas constant, *T* is temperature, 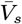 is the partial molar volume of the CPA, and 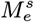 and 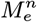 denote the external permeating and nonpermeating solute concentrations, respectively. Cells were assumed spherical with an initial volume of *V*_0_ = 1500 *µ*m^3^, consistent with a mean diameter of 14*µ*m measured during trypan blue exclusion.

To our knowledge, literature values for *L*_*p*_ and *P*_*s*_ are not available for suspended human dermal fibroblasts in propylene glycol at 4◦C. We therefore adopted parameter values reported for adherent endothelial cells under comparable conditions, which provide a reasonable surrogate(57). To extrapolate these parameters to suspended cells, we assumed a two-fold increase in accessible membrane area upon detachment from the substrate. Because transport fluxes enter as products *L*_*p*_*A* and *P*_*s*_*A*, this assumption is equivalent to using effective permeabilities 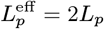 and 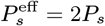 in the spherical-cell model, yielding 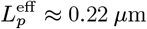 min^−1^ atm^−1^ and 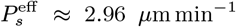 for *V*_0_ = 1500 *µ*m^3^. These parameters were used for all transport simulations shown in Figure 6.

### Warming rate simulations

Simulations of droplet warming were carried out using COMSOL Multiphysics software with the Conjugate Heat Transfer module in solids and fluids, as outlined in our previous work (29, 58). Two droplet diameters were studied, 100 µm and 200 µm, corresponding to orifice diameters of 50 µm and 100 µm, respectively. Droplet temperature-dependent thermophysical properties, including density, specific heat capacity, and thermal conductivity, were assumed to be those of 2 M propylene glycol (59).

Heat transfer within the droplet was modeled by transient heat conduction:

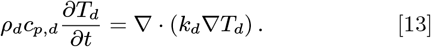

The surrounding fluid was modeled using the coupled transient energy and incompressible flow equations:

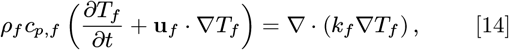

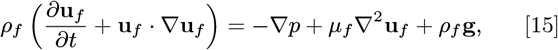

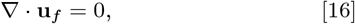

where *T*_*f*_, **u**_*f*_, *p, ρ*_*f*_, *c*_*p,f*_, *k*_*f*_, and *µ*_*f*_ are the temperature, velocity, pressure, density, specific heat capacity, thermal conductivity, and dynamic viscosity of the surrounding fluid, respectively.

Free convection was incorporated using the Boussinesq approximation:

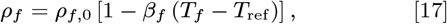

where *ρ*_*f*,0_ is the reference fluid density, *β*_*f*_ is the thermal expansion coefficient, and *T*_ref_ is the reference temperature. At the droplet–fluid interface, continuity of temperature and heat flux was imposed:

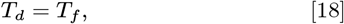

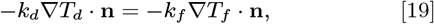

where **n** is the unit normal vector pointing from the droplet into the surrounding fluid. Thus, heat transfer from the external fluid to the droplet was determined directly from the coupled conduction and free-convection solution rather than through a prescribed convective heat-transfer coefficient due to the uncertainties involved in estimating the heat transfer coefficient. In reality, fluid motion in the rewarming bath around the droplet would aid in heat transfer, and the transition period from air to liquid involved during droplet submersion would decrease effective heat transfer rates. Proper simulation of this transition period is beyond the scope of this paper, and the rates involved here are treated as a best estimate of warming rates.

## Supporting Information

The Supporting Information is attached as a separate file.

## ACKNOWLEDGMENTS

The authors thank Dr. Bat-Erdene Namsrai and Diane K. Tobolt from Dr. Erik Finger’s laboratory for assistance with sourcing blood samples and performing hemolysis measurements using a UV spectrophotometer. We also thank Dr. Charles Elder for assistance in developing protocols for isolating and storing red blood cells. We thank NSF REU summer students Keiron Best, Ashley Everette, and Faith Neely for labeling the images used to train the computer vision algorithm, as well as high school student Warren Wang for his assistance. This work was supported by National Science Foundation Award 1941543 through the NSF Engineering Research Center for Advanced Technologies for the Preservation of Biological Systems (ATP-Bio).

## Notes

### Competing Interest Statement

The authors have declared no competing interest.

