## Supplemental Information for "Cryoaerosolization Enables Scalable Vitrification-Based Cell Cryopreservation"

PNAS

#### This PDF file includes:

Figs. S1 to S4
Tables S1 to S2
SI References

### 1. Machine Learning Training

Figure S1 compares the droplet number size distributions predicted by the machine learning model with the ground-truth measurements. In Fig. S1(a), the predicted and ground-truth histograms show close agreement. The peak of the ground-truth distribution occurs at approximately 110 $\mu$m, while the predicted distribution peaks near 115 $\mu$m, corresponding to a relative error of less than 5%. The overall distribution shape is also well captured, with similar widths and tail behavior between the predicted and ground-truth data. The slight shift in peak diameter and minor differences in the distribution tails are likely due to challenges in detecting droplet edges when droplets overlap or have poorly defined boundaries in the microscope images.

The cumulative distribution function (CDF) comparison in Fig. S1(b) provides additional insight into model performance across the full size range. The ground-truth CDF rises sharply near 100 $\mu$m, indicating that a large fraction of droplets falls within a narrow size range. The predicted CDF increases more gradually, suggesting a slight overprediction of smaller droplets and underprediction of larger droplets. Despite these differences, the predicted and ground-truth CDFs converge at both the lower and upper size limits, indicating that the model captures the overall droplet size distribution present in the dataset.

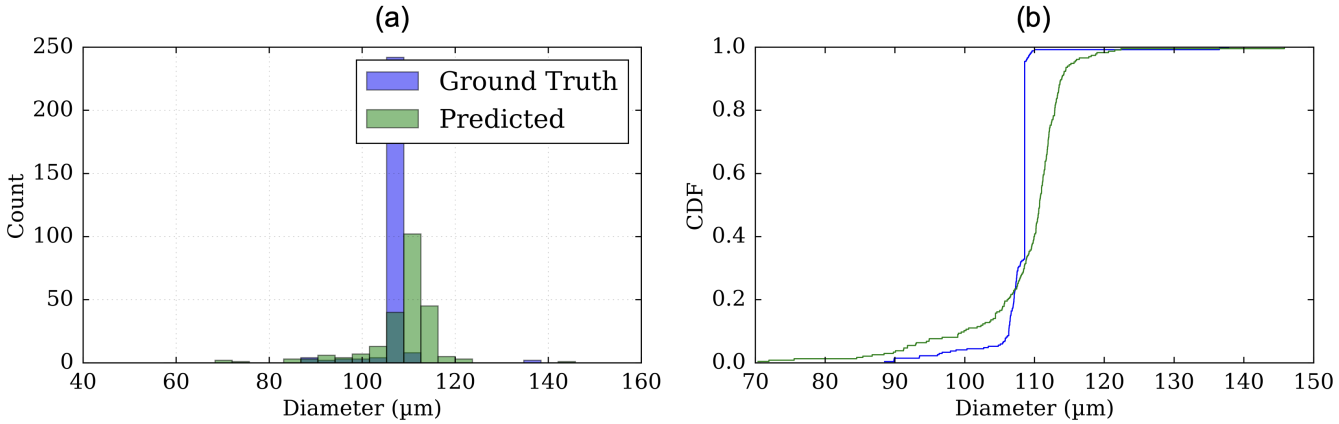


**Fig. S1.** (a) Predicted droplet size distribution compared with the ground-truth distribution. (b) Comparison of the cumulative distribution function (CDF) of droplet diameter for the ground-truth and predicted values.

The training convergence behavior of the machine learning model is shown in Fig. S2. Figure S2(a) shows the total binary cross-entropy (BCE) loss as a function of training epoch. The loss decreases monotonically and plateaus after approximately 60–70 epochs, indicating convergence of the training process. The smooth decrease in loss without substantial oscillations suggests stable training dynamics.

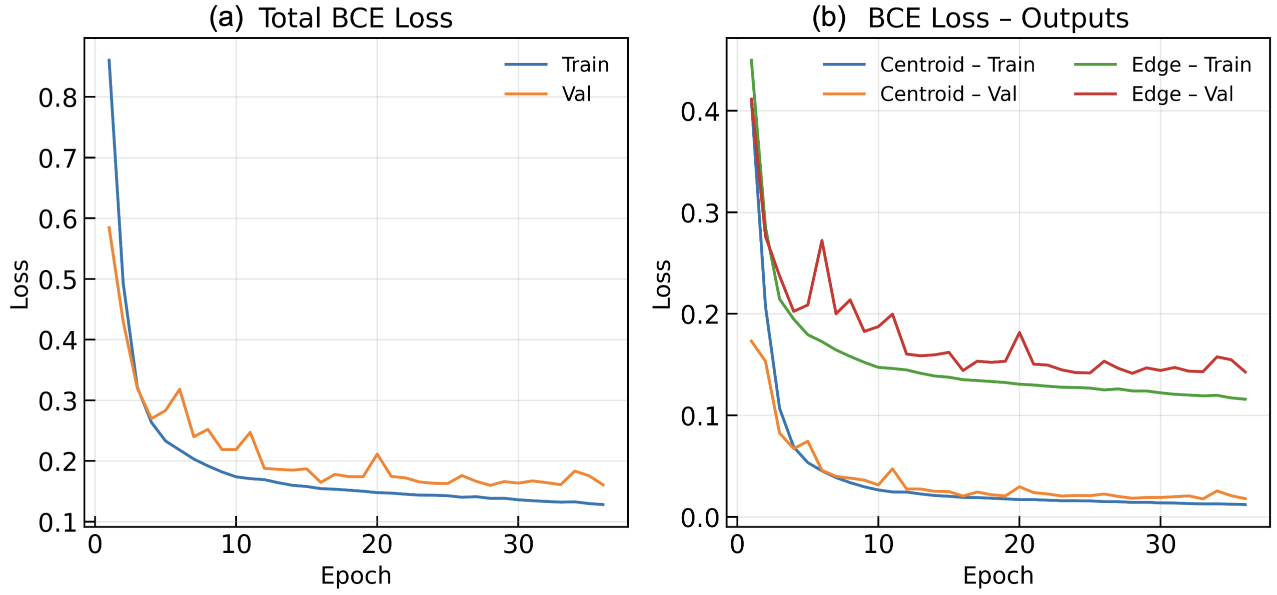


**Fig. S2.** (a) Variation of the total binary cross-entropy (BCE) loss with epoch. (b) Convergence of the centroid and edge prediction losses with epoch.

Figure S2(b) shows the individual loss components for centroid and edge prediction. The centroid loss decreases rapidly during the initial training phase and reaches a low plateau within the first 20–30 epochs. This faster convergence is expected because centroid detection requires identifying a localized region for each droplet. In contrast, the edge loss decreases more gradually and requires more epochs to converge, reflecting the greater complexity of resolving droplet boundaries across many pixels. These different convergence rates highlight the multi-scale nature of the segmentation task, in which the model learns both coarse droplet localization through centroid detection and fine-scale boundary information through edge detection.

Overall, the small discrepancy between the predicted and ground-truth size distributions is expected to have minimal effect on the final analysis because droplet size distributions were obtained by averaging over multiple images. As a result, the minor systematic differences observed in Fig. S1 are reduced when processing large numbers of droplets, yielding size distributions that are representative of the experimental conditions.

### 2. Intensity Variation

To determine an intensity threshold for classifying vitrified and frozen droplets, we recorded the temporal variation in image intensity for representative droplets. Droplets were collected and imaged, and two representative droplets were selected: one initially vitrified and one frozen. A small region of interest was drawn within each droplet, and the mean intensity within this region was recorded over a 10 min period, as shown in the intensity time-series data in Fig. S3. Vitrified droplets appeared optically transparent and exhibited lower intensity values, whereas frozen droplets scattered more light and produced higher measured intensities. Over time, initially vitrified droplets crystallized, producing a sudden increase in intensity. We also observed a similar jump in the intensity of frozen droplets due to the frost formation. Based on these measurements, an intensity threshold of 60 was selected to distinguish vitrified from frozen droplets during image classification. Optical transparency was previously validated as a surrogate for vitrification in our work comparing transparent and opaque propylene glycol droplets with crystallization measurements obtained by X-ray diffraction.(1)

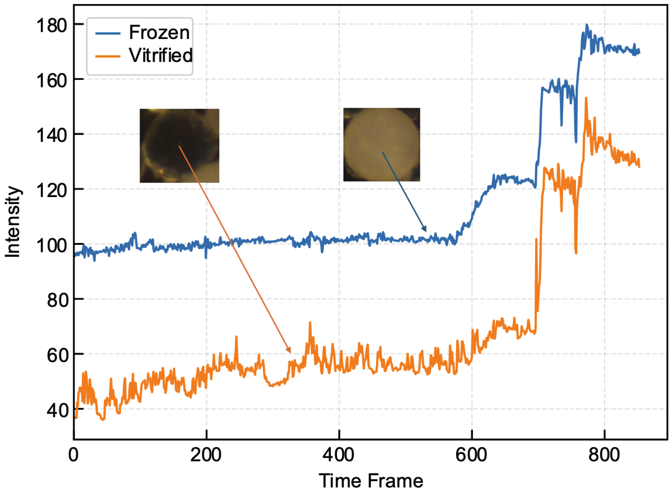


**Fig. S3.** Intensity time series data for vitrified (orange curve) and frozen droplets (blue curve). The vitrified droplets have lower intensity values than the frozen droplets.

### 3. VOAG parameters

For characterization, we varied the orifice size from 50 to 150 $\mu$m. The flow rates, frequencies, and dispersion air flow rates used for each condition are summarized in the following table.

**Table S1. Experimental parameters used with VOAG setup**

| Orifice Size ($\mu$m) | Liquid Flow Rate (mL min$^{-1}$) | Frequency (kHz) | Dispersion Air (L min$^{-1}$) |
|---|---|---|---|
| 50 | 0.6 | 17.25 | 1.5 |
| 100 | 2.2 | 5.00 | 2.5 |
| 150 | 3.8 | 2.25 | 5.0 |

### 4. Size dependence of crystallized fraction

The dependence of crystallized fraction on droplet size was evaluated as a function of ethanol concentration. At low ethanol concentrations, the crystallized fraction approached unity, indicating that nearly all droplets crystallized. As ethanol concentration increased, the crystallized fraction decreased toward zero, corresponding to vitrification of the droplet population. Larger droplets required higher ethanol concentrations to achieve the same vitrification outcome, consistent with their greater thermal mass and longer characteristic cooling times.

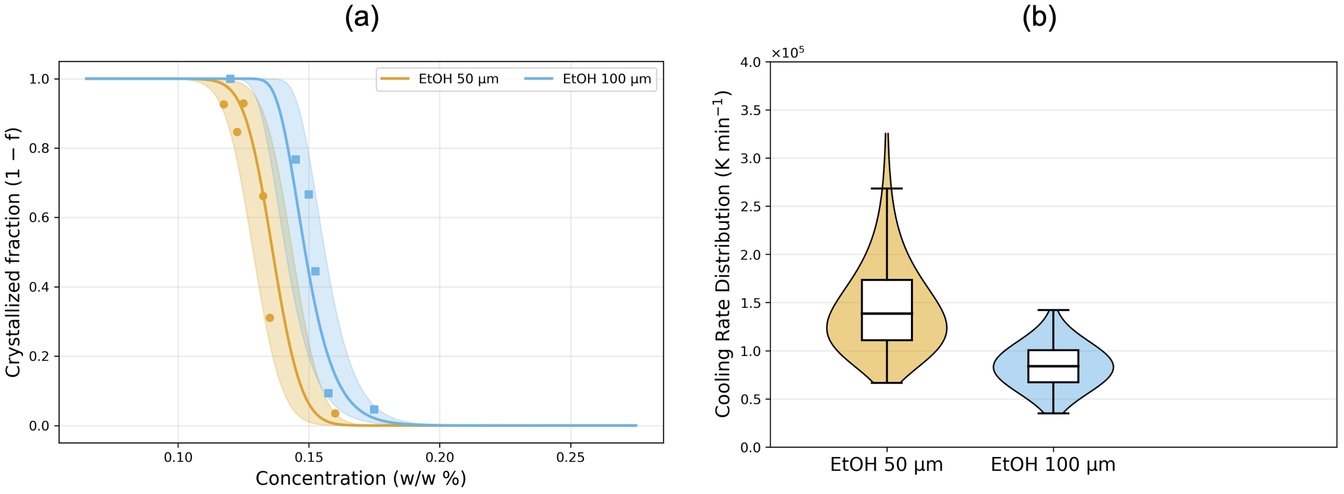


**Fig. S4.** (a) Crystallized fraction as a function of ethanol concentration (w/w %). (b) Cooling-rate distributions for different droplet sizes.

The corresponding cooling-rate distributions are shown in Fig. S4(b). The mean cooling rate was higher for 50 $\mu$m droplets than for 100 $\mu$m droplets, reflecting the higher surface-area-to-volume ratio of smaller droplets.

### 5. Mathematical model to predict droplet cooling rates

We developed a mathematical model to estimate the cooling rates of aerosolized droplets during cryoaerosolization. The model accounts for two primary heat-transfer mechanisms: convective heat transfer to the surrounding cold nitrogen gas and latent heat removal associated with collisions between aerosolized droplets and entrained liquid nitrogen droplets. For a spherical aerosolized droplet of mass $m_D$, specific heat capacity $c_p$, surface area $A_D$, and temperature $T$, the lumped energy balance is

$$m_D c_p \frac{dT}{dt} = -hA_D\left(T - T_{LN_2}\right) - \eta_{lat}\beta N_{LN_2}\Delta h_{vap} m_{LN_2}, \qquad \text{[S1]}$$

where $h$ is the gas-side convective heat-transfer coefficient, $T_{LN_2}$ is the liquid nitrogen temperature, and $A_D = \pi D^2$ for a spherical droplet of diameter $D$. The first term on the right-hand side represents convective cooling to the surrounding nitrogen gas. The second term represents latent heat removal during collisions with $LN_2$ droplets, where $\beta$ is the collision kernel between aerosolized droplets and $LN_2$ droplets, $N_{LN_2}$ is the $LN_2$ droplet number concentration, $m_{LN_2}$ is the mass of a single $LN_2$ droplet, $\Delta h_{vap}$ is the latent heat of vaporization of liquid nitrogen, and $\eta_{lat}$ is an effective latent-transfer efficiency. The temperature derivative $dT/dt$ gives the rate of temperature change, while the cooling-rate magnitude is $-dT/dt$.

**Non-Dimensionalization.** To simplify the analysis, Eq. (S1) was nondimensionalized using the temperature scale $\Delta T_* = T_i - T_{LN_2}$ and the convective cooling time scale $m_D c_p/(hA_D)$. The dimensionless temperature and time were defined as

$$\theta^* = \frac{T - T_{LN_2}}{\Delta T_*}, \qquad \tau = \frac{hA_D}{m_D c_p} t. \qquad \text{[S2]}$$

For a spherical droplet of diameter $D$, $A_D = \pi D^2$ and $m_D = (\pi/6)\rho D^3$, giving

$$\tau = \frac{6ht}{\rho D c_p}, \qquad \text{[S3]}$$

where $\rho$ is the droplet density.

Substituting these definitions into Eq. (S1) gives

$$\frac{d\theta^*}{d\tau} = -\theta^* - \frac{\eta_{lat}\beta N_{LN_2}\Delta h_{vap} m_{LN_2}}{hA_D\Delta T_*}. \qquad \text{[S4]}$$

We define the dimensionless parameter $\Phi$ as

$$\Phi = \frac{\eta_{lat}\beta N_{LN_2}\Delta h_{vap} m_{LN_2}}{h A_D \Delta T_*}, \tag{S5}$$

which represents the ratio of latent heat removal by $LN_2$ droplet impacts to gas-side convective heat removal. Eq. (S4) can therefore be written in compact form as

$$\frac{d\theta^*}{d\tau} = -\theta^* - \Phi. \tag{S6}$$

When $\Phi = 0$, the model reduces to the standard lumped-capacitance convection model. Larger values of $\Phi$ indicate a greater contribution from $LN_2$ droplet impacts. Because the latent term is treated as an average heat-removal rate, this model is used to estimate characteristic cooling rates over the relevant early cooling interval rather than to predict the full temperature history to equilibrium.

**Collision Kernel $\beta$.** The collision kernel $\beta$ describes the effective volume swept out per unit time by an aerosolized droplet as it moves relative to entrained $LN_2$ droplets. Multiplying this kernel by the $LN_2$ droplet number density gives the average collision frequency experienced by a single aerosolized droplet. For an aerosolized droplet of diameter $D$ and an $LN_2$ droplet of diameter $D_{LN_2}$, we estimated the collision kernel as the product of the collision cross section and the mean relative velocity,

$$\beta(D, D_{LN_2}) = \eta_c \frac{\pi}{4}\left(D + D_{LN_2}\right)^2 \overline{U}_{rel}, \tag{S7}$$

where $\overline{U}_{rel}$ is the mean relative velocity between the aerosolized droplets and the entrained $LN_2$ droplets, and $\eta_c$ is an effective collision efficiency. The factor of $\pi/4$ appears because $D$ and $D_{LN_2}$ are diameters rather than radii. Thus, $\beta$ has units of volume per time, and $\beta N_{LN_2}$ has units of inverse time, representing the effective collision frequency for an individual aerosolized droplet.

The latent heat-removal term assumes that collisions between aerosolized droplets and $LN_2$ droplets remove heat through vaporization of liquid nitrogen at the droplet surface. However, not every $LN_2$ droplet impact is expected to result in complete vaporization using heat transferred only from the aerosolized droplet. We therefore introduced an effective latent-transfer efficiency, $\eta_{lat}$, to account for incomplete vaporization, imperfect thermal coupling, and uncertainty in the fraction of $LN_2$ droplet mass that contributes directly to heat removal from the aerosolized droplet. For monodisperse $LN_2$ droplets, the resulting latent heat-removal rate is

$$\dot{q}_{lat} = \eta_{lat}\Delta h_{vap}\beta N_{LN_2} m_{LN_2}, \tag{S8}$$

where $\Delta h_{vap}$ is the latent heat of vaporization of liquid nitrogen, $m_{LN_2}$ is the mass of a single $LN_2$ droplet, and $N_{LN_2}$ is the $LN_2$ droplet number concentration.

**Size Distribution of Liquid Nitrogen Droplets.** The preceding expression assumes monodisperse $LN_2$ droplets. To account for a distribution of $LN_2$ droplet sizes, let $n(D_{LN_2})$ denote the $LN_2$ droplet number distribution, such that $n(D_{LN_2})\,dD_{LN_2}$ is the number density of droplets with diameters between $D_{LN_2}$ and $D_{LN_2} + dD_{LN_2}$. The latent heat-removal term can then be written as

$$\dot{q}_{lat} = \eta_{lat}\Delta h_{vap}\int_0^\infty \beta(D, D_{LN_2}) m_{LN_2}(D_{LN_2}) n(D_{LN_2})\,dD_{LN_2}. \tag{S9}$$

Using

$$m_{LN_2}(D_{LN_2}) = \frac{\pi}{6}\rho_{LN_2} D_{LN_2}^3, \tag{S10}$$

the dimensionless latent-to-convective heat-removal ratio becomes

$$\Phi = \frac{\eta_{lat}\rho_{LN_2}\Delta h_{vap}}{6hD^2\Delta T_*}\int_0^\infty \beta(D, D_{LN_2}) D_{LN_2}^3 n(D_{LN_2})\,dD_{LN_2}, \tag{S11}$$

where $h$ is the gas-side convective heat-transfer coefficient, $D$ is the aerosolized droplet diameter, and $\Delta T_* = T_i - T_{LN_2}$ is the initial temperature difference. The collision efficiency $\eta_c$ and latent-transfer efficiency $\eta_{lat}$ both reduce the effective latent heat removed per geometric collision. Since they enter the model only through their product, we define a combined effective efficiency,

$$\eta_{\text{eff}} = \eta_c\eta_{lat}. \tag{S12}$$

Substituting the collision kernel from Eq. (S7) gives

$$\Phi = \frac{\eta_{\text{eff}}\pi\rho_{LN_2}\overline{U}_{rel}\Delta h_{vap}}{24hD^2\Delta T_*}\left[D^2M_3 + 2DM_4 + M_5\right] \tag{S13}$$

where the moments of the $LN_2$ droplet size distribution are defined as

$$M_x = \int_0^\infty D_{LN_2}^x n(D_{LN_2})\,dD_{LN_2}. \tag{S14}$$

This form shows that latent heat removal increases with $LN_2$ droplet mass concentration, relative velocity, and the size-dependent collision cross section. Larger values of $\Phi$ indicate a greater contribution from $LN_2$ droplet impacts relative to gas-side convection.

**Model Parameters.** The parameters shown in Table S2 were used in the heat-transfer model calculations displayed in the main text, Fig. 1.

**Table S2. Parameters used in the heat-transfer model for cooling-rate predictions.**

| Parameter | Symbol | Value |
|---|---|---|
| Liquid nitrogen density | $\rho_{LN_2}$ | $807\,\mathrm{kg\,m^{-3}}$ |
| Latent heat of vaporization of $LN_2$ | $\Delta h_{vap}$ | $2 \times 10^5\ \mathrm{J\,kg^{-1}}$ |
| Initial temperature difference | $\Delta T_*$ | $196\,\mathrm{K}$ |
| $LN_2$ droplet geometric mean diameter | $D_{g,LN_2}$ | $500\,\mathrm{\mu m}$ |
| $LN_2$ droplet geometric standard deviation | $\sigma_{g,LN_2}$ | 1.8 |
| Mean droplet–droplet relative velocity | $\overline{U}_{rel}$ | $5.0\,\mathrm{m\,s^{-1}}$ |
| Gas–droplet relative velocity | $U_g$ | $0.02\,\mathrm{m\,s^{-1}}$ |
| VOAG frequency | $f$ | $5000\,\mathrm{Hz}$ |
| Aerosolized droplet density | $\rho_D$ | $1000\,\mathrm{kg\,m^{-3}}$ |
| Gas density | $\rho_g$ | $1.2\,\mathrm{kg\,m^{-3}}$ |
| Gas thermal conductivity | $k_g$ | $0.026\,\mathrm{W\,m^{-1}\,K^{-1}}$ |
| Gas dynamic viscosity | $\mu_g$ | $1.76 \times 10^{-5}\,\mathrm{Pa\,s}$ |
| Gas Prandtl number | $Pr$ | 0.72 |
| $LN_2$ droplet number-density range | $N_{LN_2}$ | $1 \times 10^2$ to $2 \times 10^5\ \mathrm{m^{-3}}$ |
| Combined effective efficiency | $\eta_{\mathrm{eff}}$ | 0.01–1 |

$LN_2$ droplets were assumed to follow a lognormal size distribution with the geometric mean diameter and geometric standard deviation listed in Table S2. The heat-transfer coefficient $h$ was calculated using the Ranz–Marshall correlation (2) for forced convection around a sphere,

$$Nu = 2 + 0.6Re^{1/2}Pr^{1/3}, \tag{S15}$$

where $Nu = hD/k_g$ is the Nusselt number and

$$Re = \frac{\rho_g U_g D}{\mu_g} \tag{S16}$$

is the gas-phase Reynolds number based on the aerosolized droplet diameter $D$. Here, $\rho_g$, $k_g$, $\mu_g$, and $Pr$ are the gas density, thermal conductivity, dynamic viscosity, and Prandtl number, respectively, and $U_g$ is the relative velocity between the aerosolized droplet and the surrounding gas.

### 6. Live/Dead Cell Viability Assay

Following CPA/aerosol spray exposure and rewarming, hiPSCs were collected in the mTeSR Plus culture medium and counted using a hemocytometer. Cells were seeded at a density of $2 \times 10^5$ cells per well in Matrigel-coated 12-well plates containing mTeSR Plus medium supplemented with 10 $\mu$M ROCK inhibitor (Selleck Chemicals, #S1049) and 1× Clone R2 (STEMCELL Technologies, #100-0691). Cells were cultured for 24 hours to allow attachment and recovery. Cell viability was then assessed using the Live/Dead Viability/Cytotoxicity Kit (Thermo Fisher Scientific, #L3224) according to the manufacturer's instructions. Briefly, cells were incubated with the Live/Dead staining solution containing Calcein AM and Ethidium Homodimer-1. Following staining, the dye-containing medium was removed, cells were washed once with 1× DPBS, and fresh culture medium was added. Fluorescence images were acquired using an EVOS M5000 fluorescence microscope (Invitrogen, USA). Live cells were stained with Calcein AM (green fluorescence) and dead cells were stained with Ethidium Homodimer-1 (red fluorescence) were quantified using AutoCount, an ImageJ-based automated cell-counting tool(3). Cell viability was calculated using the following equation.

$$\text{Cell Viability } (\%) = 100 \times \frac{\text{Number of Live Cells}}{\text{Number of Live Cells} + \text{Number of Dead Cells}} \tag{S17}$$

The viability of each treatment group was subsequently compared with that of the untreated control group.

### 7. Colony Formation Assay

The colony formation assay was performed as previously described with minor modifications(4). Following treatment, hiPSCs were harvested, counted, and seeded at a density of $1 \times 10^4$ cells per well in Matrigel-coated six-well plates. Cells were cultured in mTeSR Plus medium supplemented with 10 $\mu$M ROCK inhibitor and 1× Clone R2. Fresh mTeSR Plus medium was replenished every three days, and cells were cultured for 5 days to allow recovery and colony formation. After 5 days of hiPSC culture, colonies were fixed with a methanol: acetic acid solution (1:1, v/v) for 30 minutes and washed three times with 1× phosphate-buffered saline (PBS). Fixed colonies were stained with 0.5% crystal violet (Sigma-Aldrich, #548629) prepared in methanol for 30 minutes. Excess stain was removed by gently washing the wells with Milli-Q water, and plates were air-dried.

The stained hiPSC colonies were counted manually, and colony formation efficiency was calculated relative to the untreated control group using the following equation.

$$\text{Colony Formation (\% of Control)} = 100 \times \frac{\text{Number of Colonies in Treatment Group}}{\text{Number of Colonies in Control Group}} \quad \text{[S18]}$$

The percentage of colonies formed in each treatment group was subsequently compared with that of the control group to assess post-treatment recovery and proliferative capacity.
